# An abundant merozoite surface protein of *Plasmodium falciparum* modulates susceptibility to inhibitory antibodies

**DOI:** 10.1101/2025.05.13.653866

**Authors:** Isabelle G. Henshall, Jill Chmielewski, Dimuthu Angage, Ornella Romeo, Keng Heng Lai, Kaitlin R. Turland, Nicki Badii, Michael Foley, Robin F. Anders, James Beeson, Danny W Wilson

## Abstract

Malaria merozoite surface proteins (MSPs), are thought to have important roles in red blood cell (RBC) invasion and their exposure on the parasite surface makes them attractive vaccine candidates. However, their role in invasion has not been directly demonstrated and their biological functions are unknown. One of the most abundant proteins is *Pf*MSP2, which is likely an ancestral protein that has been maintained in the *Plasmodium falciparum* lineage and is a focus of vaccine development, whose function remains unknown. Using CRISPR-Cas9 gene-editing, we removed *Pf*MSP2 from two different *P. falciparum* lines with no impact on parasite replication or phenotype *in vitro*, demonstrating that it is not essential for RBC invasion. However, loss of *Pf*MSP2 led to increased inhibitory potency of antibodies targeting other merozoite proteins involved in invasion, particularly *Pf*AMA1. In a solid-phase model, increasing concentrations of *Pf*MSP2 protein reduced binding of different antibodies against *Pf*AMA1 in a dose dependent manner. These data suggest that *Pf*MSP2 can modulate the susceptibility of merozoites to protective inhibitory antibodies. The results of this study change our understanding of the potential functions of *Pf*MSP2 and establishes a new concept in malaria where a surface protein can reduce the protective efficacy of antibodies targeting a different antigen. These findings have important implications for understanding malaria immunity and informing vaccine development.

## Introduction

*Plasmodium falciparum* malaria remains a global health challenge with over half a million deaths annually (World Health Organization, 2024). Worryingly, increasing parasite resistance to frontline antimalarials and mosquito insecticide resistance are decreasing the effectiveness of key control measures (World Health Organization, 2024). Development of the RTS,S and R21 vaccines targeting the *P. falciparum* circumsporozoite protein is a step forward for malaria vaccine development, however further improvements in vaccine efficacy and longevity are needed to protect those most at risk – children under 5 years – and enable malaria elimination (Beeson et al., 2019; Datoo et al., 2024; RTSS Clinical Trials Partnership, 2015). Understanding the function of vaccine targets remains important for prioritisation and optimisation of candidates in the development pipeline. Merozoites, the parasite stage that invades host red blood cells (RBCs), have long been targeted for vaccine development since successful blocking of merozoite invasion would prevent the repeated cycles of blood-stage parasite multiplication and associated disease. While much work has been done to understand the processes and steps required for merozoite invasion, only a few merozoite proteins have successfully had functions defined. To date, most vaccines based on merozoite antigens have demonstrated limited efficacy (Beeson et al., 2016) at least in part due to the extensive polymorphisms exhibited by many of these antigens.

The surface of the merozoite is covered in a fibril coat which is comprised of multiple proteins, broadly known as merozoite surface proteins (MSPs). These proteins have been proposed to mediate weak initial merozoite RBC contact (Bannister et al., 1986; Beeson et al., 2016; Cowman et al., 2017; Weiss et al., 2015) and in *P. falciparum* are dominated by GPI-anchored proteins, the most abundant being *Pf*MSP1 and 2 (Gilson et al., 2006a). *Pf*MSP1 has been suggested to have a role in merozoite attachment to the RBC (Li et al., 2004), but recent evidence instead suggests that it has a role in merozoite rupture from schizonts and cellular interaction studies using optical tweezers do not support a substantial role for this protein in RBC binding (Das et al., 2015; Kals et al., 2024), although it may have another function in invasion. These recent insights into *Pf*MSP1 function, the best studied MSP, highlight what little is known of MSPs and their roles in parasite survival.

*P. falciparum* merozoite surface protein 2 (*Pf*MSP2), an antigen reported to be refractory to gene knock-out in *P. falciparum* (Sanders et al., 2006) but that has also been reported to be dispensable in a piggyBac mutagenesis study (Zhang et al., 2018), has been of long-term interest as a vaccine candidate. *Pf*MSP2 is a ∼28 kDa intrinsically disordered protein that has conserved N- and C- termini, along with a central variable region that defines two main allelic groups, 3D7-like and FC27-like (Adda et al., 2012). While *Pf*MSP2 was theorised to have a mechanical role in the early steps of invasion (Anders et al., 2010), there is minimal supporting evidence for this, in part due to the lack of tools to study *Pf*MSP2 function. Studies with recombinant proteins suggest that it is likely *Pf*MSP2 interacts with lipids and forms complexes with itself on the merozoite surface (Adda et al., 2009; Lu et al., 2019; Zhang et al., 2012). *Pf*MSP2 peptides have also been reported to bind the RBC and inhibit *P. falciparum* growth, which would support *Pf*MSP2 mediating merozoite-RBC interactions (Ocampo et al., 2003). Unlike the majority of the merozoite surface coat proteins, *Pf*MSP2 is not present in other malaria parasites that infect humans and has been postulated to have evolved specifically in the *Laverania* subgenus of *Plasmodium*, which includes *P. falciparum* (Black et al., 2002).

*Pf*MSP2 has been trialled in a vaccine (Combination B) combined with fragments of *Pf*MSP1 and *Pf*RESA. A Phase 1/2b trial of this vaccine showed efficacy in reducing the parasite burden in children, which was specific to the *Pf*MSP2 3D7-like allele used in the vaccine formulation (Genton et al., 2003). Vaccine-induced antibodies from clinical trials or experiments in animals have little or no inhibitory activity in standard growth inhibition assays (Boyle et al., 2015; McCarthy et al., 2011), but did show functional activity through the recruitment of complement, promoting opsonic phagocytosis and antibody-dependent cellular inhibition (Boyle et al., 2015; Feng et al., 2018; McCarthy et al., 2011). Naturally acquired antibodies are also known to target *Pf*MSP2 and have been linked to protection (Fowkes et al., 2010; Osier et al., 2010; Stanisic et al., 2009; Zerebinski et al., 2024). Studies have highlighted that it is the Fc-mediated functional activities of these naturally acquired *Pf*MSP2 antibodies, as opposed to direct blocking of *Pf*MSP2 protein function, that correlates best with protection (Boyle et al., 2015, 2014; Osier et al., 2014; Reiling et al., 2019). While *Pf*MSP2 is a key target of naturally acquired malaria immunity, little is known about the function of *Pf*MSP2, hindering its advancement as a potential vaccine candidate.

Here we take a reverse genetics approach to assess the function of *Pf*MSP2, quantifying the impacts of *Pf*MSP2 deletion on parasite invasion and phenotype. Given our findings that deletion of MSP2 did not impact invasion *in vitro*, we tested the alternative hypothesis that merozoite surface proteins may function to modulate susceptibility of merozoites to inhibitory antibodies.

## Results

### Avian malaria MSP2-like proteins are structurally similar to *Laverania* MSP2s and show that MSP2 did not arise exclusively in this clade

Genome annotations of *pfmsp2* show it lies between the merozoite surface protein *pfmsp5* (PF3D7_0206900) and *pfmsp*4 (PF3D7_0207000), and the conserved purine biosynthesis enzyme *adenylosuccinate lyase* (ADSL, PF3D7_0206700) (Figure 1A) (PlasmoDB; (Aurrecoechea et al., 2009)), with this arrangement previously thought to be restricted to the *Laverania* clade which includes parasites that infect great apes and *P. falciparum* (Black et al., 2002; Ferreira et al., 2015). As described by Escalante et al. (2022), we found the genomes of the two annotated avian *Plasmodium* spp. available on PlasmoDB, *P. relictum* and *P. gallinaceum*, to also have a putative gene between ADSL and MSP5 with structural similarities to *Laverania* MSP2s. The reduced length of the *P. relictum* (PRELSG_0415300) and *P. gallinaceum* (PGAL8A_00017700) MSP2-like proteins compared to those of *Laverania* parasites is accounted for by the absence of any extended central domains of the two avian malaria parasites (Supplementary Figure 1).

**Figure 1.**
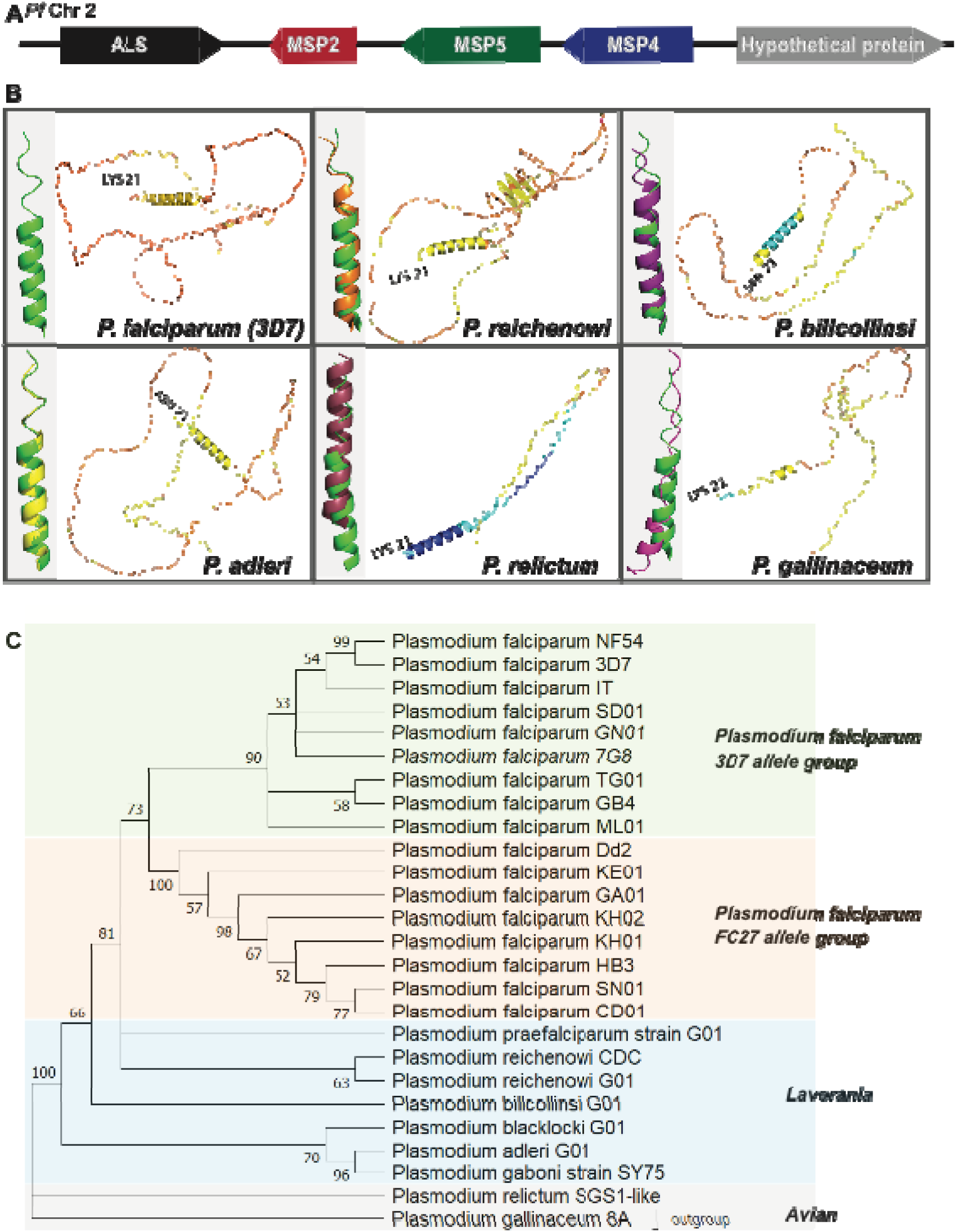
Presence and structure of MSP2 across different *Plasmodium* species. **(A)** Schematic of the gene arrangement surrounding *msp2* on *P. falciparum* chromosome 2. **(B)** AlphaFold2 structural predictions of example *Laverania* (*P. falciparum* (3D7), *P. reichenowi*, *P. billcollinsi*, *P. adleri*) and the two avian (*P. relictum*, *P. gallinaceum*) malaria species. The N-terminal signal peptide and C-terminal GPI anchor sequences were removed before the proteins structure was predicted. The most N-terminal amino acid is indicated. Colours represent the predicted local distance difference test (pLDDT) scores, with dark blue representing very high confidence (>90%), light blue high confidence (90 to >70), yellow low confidence (70 to 50) and orange very low confidence (<50). An enlarged modelled structure of the N-terminal helical region is provided for *P. falciparum* 3D7 MSP2 on the left of the panel (grey shading), and also for the other examples of *Laverania* and avian MSPs (various colours) superimposed on the structure of *P. falciparum* 3D7 MSP2 (green). **(C)** Maximum likelihood tree showing relationship of MSP2 protein sequences found in different *P. falciparum* isolates and other *Plasmodium* species. Grouping of sequences from *P. falciparum* isolates into two main allele types can be seen as well as the separation of the *Laverania* and avian malaria species into their expected groups. Tree robustness tested by bootstrap.

Given the estimated that ∼10 million years of evolution separate the avian malaria parasites *P. relictum* and *P. gallinaceum* from the *Laverania* (Böhme et al., 2018), we explored whether similarities in amino acid sequence and predicted conformation would support a shared lineage and functional properties of these putative MSP2s. Although *Pf*MSP2 is generally considered a disordered protein with limited conservation, the hydrophobic N- and C-terminal sequences that correspond to secretion signals (SP) and GPI-attachment signals, as well as short regions downstream and upstream of these signal sequences respectively, show significant conservation across *P. falciparum* MSP2s. Here we compared the sequence similarity of these conserved regions between the *Laverania* and avian MSPs and found consistently high levels of conservation for the putative SP (>89% similarity) (Supplementary Figure 1), the 24 amino acid N-terminal conserved region (78% similarity) that has a propensity for alpha-helical structure and fibril formation (Yang et al., 2010) (Supplementary Figure 2), the 50 amino acid residue C-terminal conserved region (62% similarity) (Supplementary Figure 3) and the C-terminal GPI signal sequence (>79%), including the presence of short side chain amino acids that mark predicted GPI cleavage sites (Eisenhaber et al., 1999; Gilson et al., 2006; Smythe et al., 1991) (Supplementary Figure 1). *Laverania* MSP2s contain two C-terminal cysteine residues that form an intramolecular disulphide bond (Zhang et al., 2008). This arrangement of 2 C-terminal cysteines is found in *P. gallinaceum* MSP2, but not *P. relictum* which has only one (Supplementary Figure 3). Overall, these N and C-terminal conserved regions show broad conservation in properties across the *Laverania* and avian malarias which could contribute to a conserved sub-cellular localisation.

Characteristic of *P. falciparum* MSP2s is the dimorphic central variable repeat region that are grouped as 3D7 or FC27-like amino acid sequence structures. A defining feature of the 3D7-like MSP2 family are 4 or 8-mer tandem repeats consisting of small amino acids (for 3D7: Gly, Gly, Ser, Ala) (MacRaild et al., 2015; Smythe et al., 1991). The FC27-like MSP2 family also has repeats of 12 or 32 amino acids in length, but these do not resemble those of the 3D7-like family. We found that the non-*P. falciparum Laverania* and avian malaria MSP2 sequences also contained >75% small amino acids (some of: Gly, Ser, Ala, Thr) repeats, but the exact number (4 to 8 repeats), length (3 to 7 amino acids) and the level of conservation between repeats differed between and within species (Supplementary Figure 4). Broadly speaking, these *Laverania* and avian malaria MSP2 repeats had a 3D7-like repeat structure, with no example of a FC27-like repeat structure identified even with an additional nine *Laverania* MSP2s from the NCBI experimental database compared (data not shown). This comparison highlights the surprising finding that a region of known dimorphic variability in *P. falciparum* MSP2 is likely to retain functional constraints that favour small amino acid rich repeats immediately downstream of the N-terminal conserved region.

Extensive studies on recombinant proteins of the *P. falciparum* 3D7 and FC27 variants of *Pf*MSP2 have shown this merozoite surface protein to be intrinsically disordered. Given the evidence for conservation of the primary structure of MSP2s from distantly related malaria parasites, we used AlphaFold2 predictions (Jumper et al., 2021) to examine whether there were likely to be also conformational similarities between *Laverania* MSP2s and those of *P. relictum* and *P. gallinaceum*. The MSP2s in the database of AlphaFold2-predicted structures retain their N- and C-terminal signal sequences, but as these are both absent from mature MSP2 on the merozoite surface they were deleted from the MSP2 sequences used for the AlphaFold2 predictions reported here. The predicted structures of the two avian parasite MSP2s, and other *Laverania* MSP2s, are very similar to that of *Pf*3D7 MSP2, with most of the polypetide chains lacking predicted secondary structure and, from the colour coding, have a predicted local distance test (pLDDT) consistent with that of an intrinsically disordered protein over the majority of the proteins length (Figure 1B). In contrast to the rest of the polypeptide, some α-helical structure was predicted in the conserved 20-residue N-terminal region of all the *Laverania* and avian MSP2s. NMR studies with recombinant *Pf*3D7 and *Pf*FC27 MSP2 showed this conserved N-terminal sequence to have a propensity for forming an α-helix that was stabilized in the presence of lipid mimetics (MacRaild et al., 2012). The low pLDDT values are consistent with this region in the other *Laverania* and *P. gallinaceum* MSP2 having some propensity for α-helix formation, whereas higher pLDDT values suggest the possible presence of a more stable α-helix in this region of *P. relictum* MSP2.

Having established that MSP2 is present across different *Plasmodium* lineages sharing similar predicted amino acid compositions and structural features, the phylogenetic relationship between annotated *Laverania* and putative avian malaria parasite MSP2s was examined. A phylogenetic tree was generated based on alignments of representative sequences from each species. MSP2 sequences from species belonging to the *Laverania* subgenus all grouped together with two smaller clusters, corresponding to the known *Laverania* clade A and clade B species (Figure 1C). As expected, *P. relictum* and *P. gallinaceum* MSP2 also group together. The phylogenetic tree generated based on MSP2 sequences mirrors the proposed evolution of *Plasmodium* spp. (Figure 1C). As postulated by Escalante et al. (2022) , these data suggest that MSP2 was likely present in the ancestral *Plasmodium* parasite prior to the split of the mammalian infecting *Plasmodium* spp. From those that infect birds and lizards. Given the shared amino acid sequence and predicted conformational similarities, including the high levels of intrinsically disordered sequence outside of the N- and C-terminal regions, the *P. relictum* and *P. gallinaceum* MSP2-like proteins appear likely to contain enough properties to potentially be functionally equivalent to *Laverania* MSP2s. This conservation of MSP2-like proteins across different species suggests it has an important function.

### Loss of *Pf*MSP2 does not noticeably impact growth or invasion of different *P. falciparum* lines *in vitro*

Given previous unsuccessful attempts to disrupt *pfmsp2* (Sanders et al., 2006), and its high abundance on the merozoite surface (Gilson et al., 2006), *Pf*MSP2 has been traditionally viewed as an essential *P. falciparum* protein with an essential function in merozoite invasion, although more recent piggyBac mutagenesis studies have called this understanding into question (Zhang et al., 2018). We employed CRISPR-Cas9 gene editing to determine whether the reported inability to knock-out *pfmsp2 in vitro* was a result of this protein being essential for parasite survival or because of the lower efficiency of the previously used gene-editing system (Sanders et al., 2006). Unexpectedly, we confirmed successful disruption of *pfmsp2* by replacing the coding sequence between 132 bp and 819 bp of the gene with a hDHFR drug selection cassette in the 3D7 *P. falciparum* laboratory-adapted line (Figure 2A and B), resulting in *Pf*3D7 ΔMSP2 parasites. We confirmed that *Pf*MSP2 was no longer expressed in *Pf*3D7 ΔMSP2 parasites by western blot, which detected a MSP2 band around 50 kDa in *Pf*3D7 WT parasite material but not in *Pf*3D7 ΔMSP2 parasites (Figure 2C). Similarly, immunofluorescence microscopy using anti-*Pf*MSP2 rabbit polyclonal antibodies showed the expected surface localisation of MSP2 on *Pf*3D7 WT merozoites but not on *Pf*3D7 ΔMSP2 parasites (Figure 2D). We next assessed whether knock-out of *Pf*MSP2 impacted on parasite growth over one or two cycles of development *in vitro* and found that deletion of *Pf*3D7 MSP2 did not cause any significant change in growth (Figure 2E and F).

**Figure 2.**
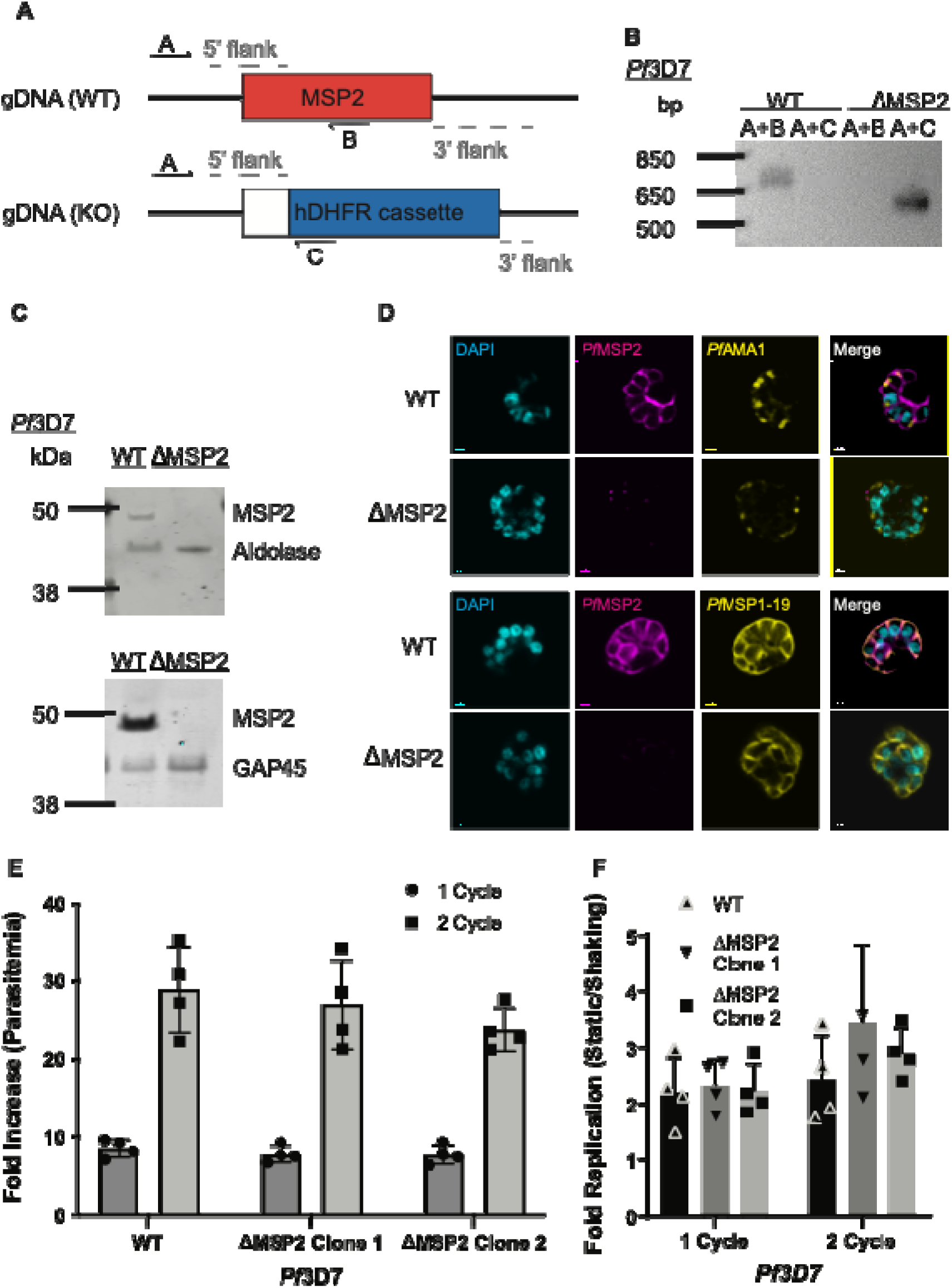
*Pf*MSP2 is not essential for growth *in vitro* of *Pf*3D7 blood stage parasites. (A-B) Schematic and agarose gel showing integration of knockout construct (band A+C) in the *msp2* gene locus and absence of the *msp2* sequence from the *Pf*3D7 ΔMSP2 Clone 1 (band A+B). **(C)** Western Blot of late schizont protein extracts confirms no *Pf*MSP2 is expressed in *Pf*3D7 ΔMSP2 Clone 1. *Pf*MSP2 detected by anti-*Pf*MSP2 2F2 3D7 mAb with *Pf*Aldolase (upper blot) or *Pf*GAP45 (lower blot) serving as loading and stage of expression controls respectively. Representative image shown. **(D)** Distribution of key merozoite surface proteins for *Pf*3D7 WT compared to *Pf*3D7 ΔMSP2 Clone 1parasites visualised by immunofluorescence. *Pf*MSP2 (magenta), the nucleus stained by DAPI (cyan) and *Pf*AMA1 (yellow, top two rows) or *Pf*MSP1-19 (yellow, bottom two rows), and the coloured merge of the preceding panels. Scale bar = 0.7 µm. Representative images shown from a minimum of 10 schizonts imaged per condition. **(E-F)** Growth of *Pf*3D7 WT compared to *Pf*3D7 ΔMSP2 Clone 1 and 2 *P. falciparum* parasites, measured as fold increase in parasitaemia, over one (48 hrs) or two (96 hrs) cycles in either standard (still- (E)) or shaking (F) conditions, with no measurable difference between parasite growth rates seen between standard or shaking conditions. Parasitaemia was determined by flow cytometry at the start and end to calculate fold increase. Graph displays mean ± S.D. of three independent experiments performed with technical triplicates. Significance determined by unpaired t-test with p< 0.05 deemed significant.

As a previous reverse genetics study in 3D7 reported that *Pf*MSP2 was essential for *P. falciparum* growth *in vitro* (Sanders et al., 2006), we investigated whether *Pf*MSP2 could also be removed from *Pf*Dd2, an isolate of *P. falciparum* that differs from 3D7 in geographical origin, RBC receptor usage and allelic type of *pfmsp2*. Using CRISPR-Cas9 we successfully knocked-out *msp2* in *Pf*Dd2 (Figure 3 A and B) and confirmed the absence of the protein in *Pf*Dd2 ΔMSP2 parasites using western blot (Figure 3C). As we found for *Pf*3D7, deletion of *Pf*Dd2 *msp2* did not result in a significant impact on parasite growth over one or two cycles of parasite replication (Figure 3D).

**Figure 3.**
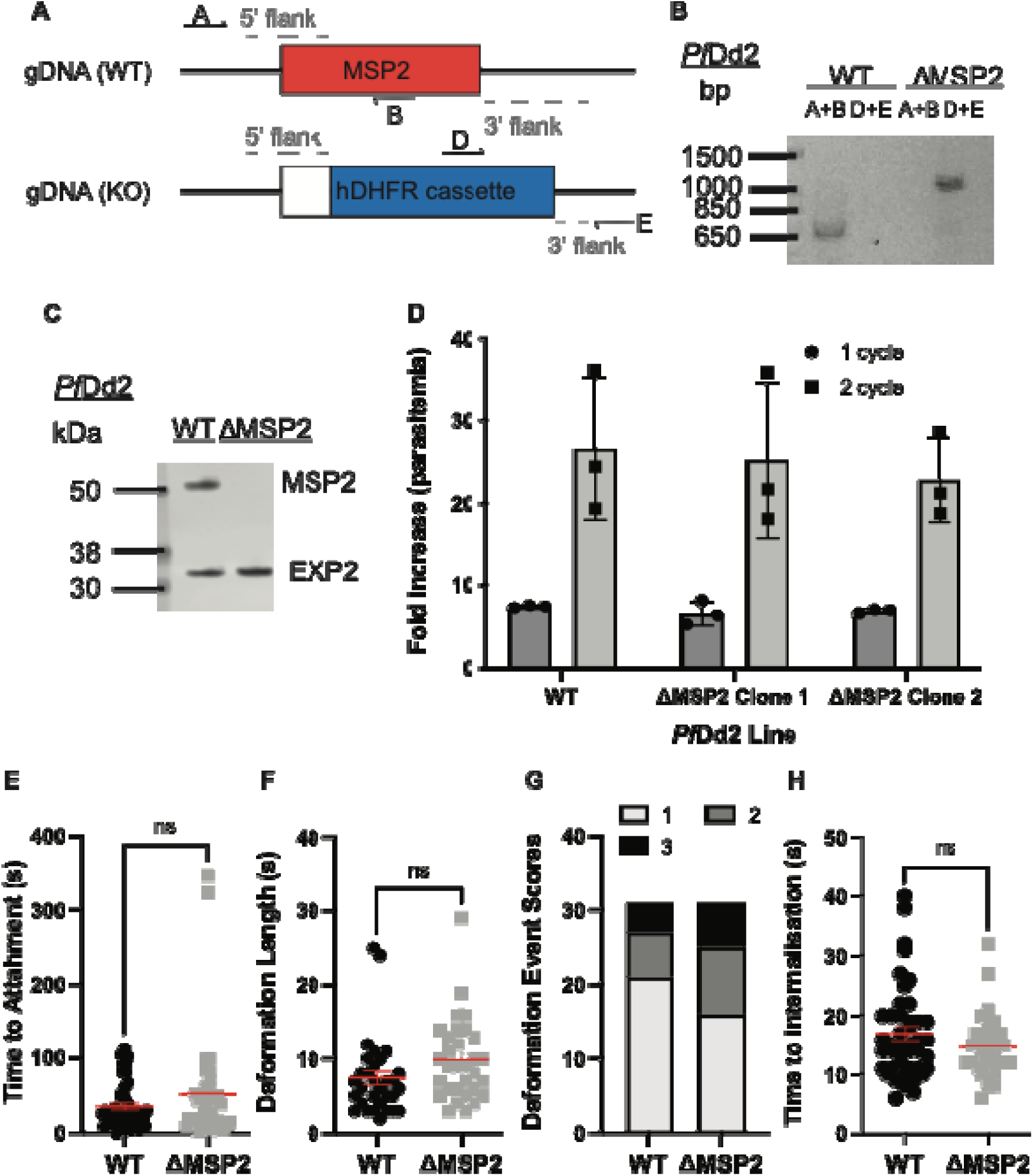
*Pf*Dd2 does not require MSP2 for asexual growth *in vitro*. (A-B) Successful integration of KO construct (schematic in A) into the *msp2* gene locus of *Pf*Dd2 ΔMSP2 Clone 1 was confirmed by PCR of genomic DNA (primers A+B amplify WT locus, primers D+E amplify integrated KO construct). **(C)** Loss of *Pf*MSP2 expression (*Pf*Dd2 ΔMSP2 Clone 1) was demonstrated by western blot of schizont protein extract with *Pf*MSP2 detected by anti-*Pf*MSP2 FC27 and anti-*Pf*EXP2 as loading control. Representative image shown. **(D)** Growth of *Pf*Dd2 WT *P. falciparum* parasites and *Pf*Dd2 ΔMSP2 Clone 1 and 2 parasites over one (48 hrs) or two (96 hrs) cycles. Parasitaemia was determined by flow cytometry at the start and end to calculate fold increase. Graph displays mean ± S.D. of three independent experiments performed with technical triplicates. **(E-H)** Key parameters of merozoite invasion were measured for both *Pf*Dd2 WT (n = 43) and *Pf*Dd2 ΔMSP2 Clone 1 (n = 35) parasites that had successfully invaded a RBC using live cell imaging of merozoite invasion. Time to merozoite attachment to RBCs **(E)**, length **(F)** and strength **(G)** of RBC deformation, and time to complete merozoite invasion **(H)** were measured by live microscopy.

Given the abundance of *Pf*MSP2 on the merozoite surface, we postulated that loss of this protein may impact on the timing of merozoite invasion of the RBC, which is not reflected in reduced growth in *in vitro* cultures. Previous studies using live-cell microscopy have reported that merozoites first contact the RBC and form weak initial interactions. The merozoite then reorientates to bring apical organelles into position to release their contents onto the RBC, which then leads to formation of the tight junction, merozoite invasion through the tight-junction, formation of the parasitophorous vacuole and resealing of the RBC membrane post-entry (Weiss et al., 2015). Given its surface localization, *Pf*MSP2 has been speculated to act in the early phase of merozoite attachment. To assess whether knock-out of *pfmsp2* impacted on the progression of invasion, we used live-cell microscopy of *Pf*Dd2 ΔMSP2 parasites and compared the timing and key steps of invasion to *Pf*Dd2 WT parasites *in vitro*. This analysis showed that, although there was a trend for *Pf*Dd2 ΔMSP2 knock-out parasites to have a higher mean time to attach to the RBC, as well as for the length and strength of RBC deformation, these trends did not reach significance. For those merozoites that did invade the RBC, on average it took less time for *Pf*Dd2 ΔMSP2 knock-out parasites to invade then *Pf*Dd2 WT, but this again did not reach significance (Figure 3 E-H). Together these data show *Pf*MSP2 is not essential for blood-stage replication *in vitro* in two *P. falciparum* laboratory isolates from different geographical regions and knock-out of *Pf*MSP2 does not seem to significantly impact parasite growth or merozoite invasion *in vitro*. However, its conservation across diverse *Plasmodium* spp. suggests it does play an important function or is advantageous for parasite survival.

Deformation scores are as defined by Weiss et al (2015), with 1 = weak deformation of the RBC membrane at the point of contact, 2 = strong deformation leading to the RBC membrane extending up the sides of the merozoite and changes in RBC membrane curvature beyond the point of contact and 3 = extreme deformation indicated by the merozoite being deeply embedded in the RBC membrane and strong deformation of the RBC well beyond the point of contact. Significance determined by unpaired t-test with p< 0.05 deemed significant.

### Impact of MSP2 gene deletion on gene expression and invasion pathway usage

Because of the close proximity of the *pfmsp*2, 4 and 5 genes on chromosome 2, similarities in their structure (e.g. GPI-anchor intrinsically disordered protein) and shared merozoite surface localization, we used qPCR to assess whether knock-out of *Pf*3D7 MSP2 impacted the expression levels of *Pf*3D7 MSP4 or MSP5. We found no change in MSP4 and MSP5 expression levels between *Pf*3D7 ΔMSP2 parasites and *Pf*3D7 WT parasites (Figure 4A). As expected, *Pf*MSP2 was not detected in the *Pf*3D7 ΔMSP2 parasites.

**Figure 4.**
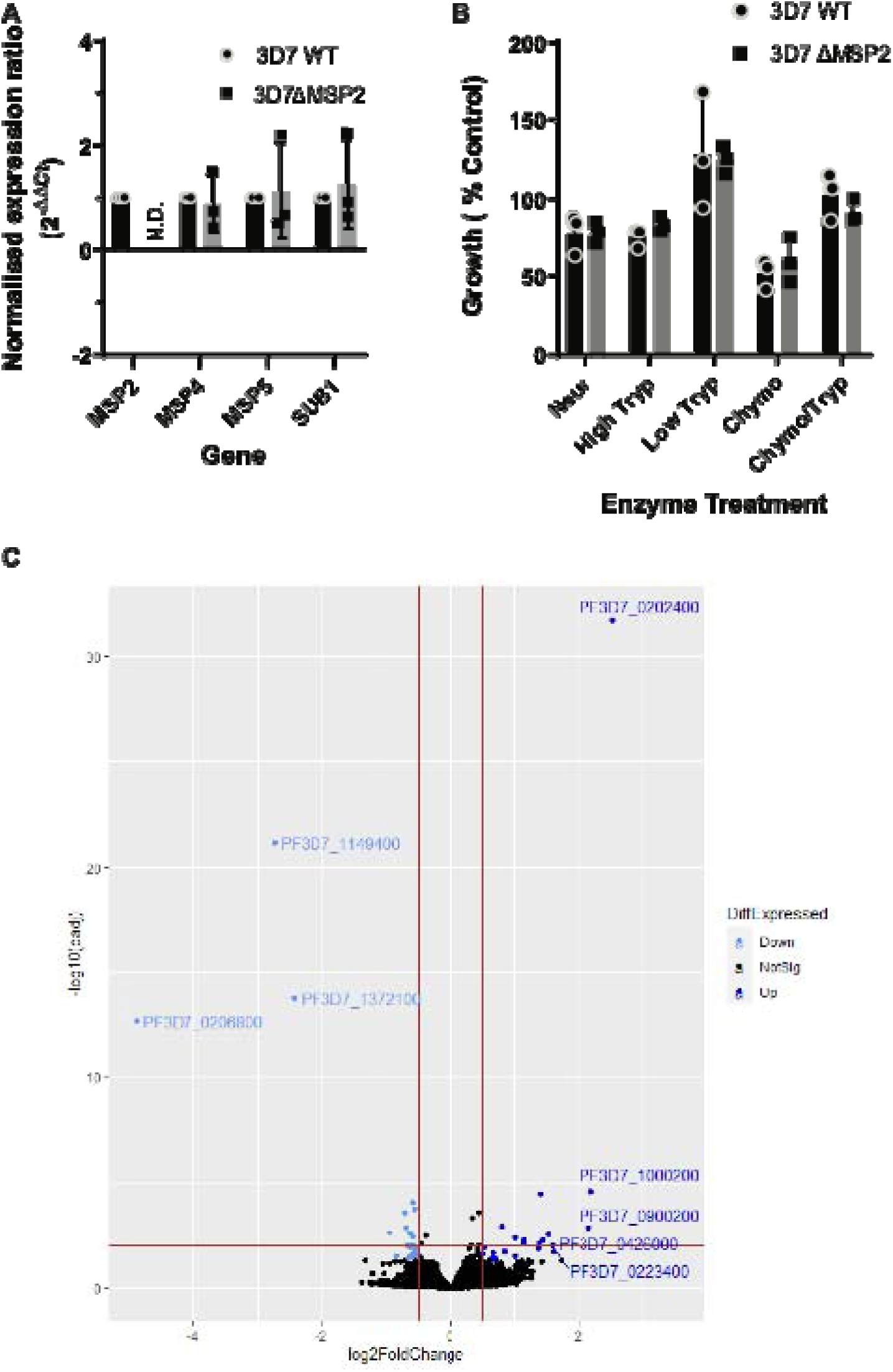
The impact of the loss of *Pf*MSP2 on expression of known merozoite invasion genes and invasion pathway utilization. **(A)** Impact of *Pf*MSP2 KO on schizont transcript abundance was assessed by qPCR for genes located in proximity to *Pfmsp*2 on chromosome 2. Changes in expression between *Pf*3D7 WT and *Pf*3D7 ΔMSP2 Clone 1 parasites was determined by qPCR relative to *pfaldolase* expression with *pfsub1* serving as a schizont stage control. Graph displays mean ± S.D. of three independent RNA harvests. **(B)** Selective enzymatic cleavage of key RBC receptors showed no difference in invasion preference between *Pf*3D7MSP2 WT and *Pf*3D7 ΔMSP2 Clone 1 parasites. Parasitaemia was determined by flow cytometry and compared to growth in non-treated control RBCs. Graph displays mean ± S.D. of three independent experiments. **(C)** Log2(fold change) for differentially expressed genes, including multigene families, between the transcriptome of *Pf*3D7 WT and *Pf*3D7 ΔMSP2 Clone 1 schizonts. Plot represents the results for one of four independent schizont RNA harvests for *Pf*3D7 WT and *Pf*3D7 ΔMSP2 parasites and red lines differentiate genes with a log2 (fold change) > 0.5 and < -0.5 with adjusted p-value < 0.01. Genes shaded blue represent those genes that were found to have an average log2 (fold change) > 0.5 (dark blue) or < -0.5 (light blue) across the four replicate samples compared. Significance determined as below p< 0.05 after correction for multiple testing.

Secreted *P. falciparum* antigens belonging to the erythrocyte binding antigen (EBA) and reticulocyte binding homologue (Rh) families are known to facilitate host-cell invasion through specific receptors on the RBC surface. Reliance on specific EBA or Rh proteins, or their cognate host-cell receptor, define different invasion pathways that can be crudely described by the sensitivity of merozoite invasion to cleavage of RBC surface proteins with trypsin, chymotrypsin and neuraminidase (Baum et al., 2005; Duraisingh et al., 2003; Orlandi et al., 1992).When we assessed whether *Pf*MSP2 knock-out impacted merozoite invasion pathway usage, we found that *Pf*3D7 ΔMSP2 parasites grew equally as well as *Pf*3D7 WT parasites in all enzyme-treated RBCs (Figure 4B).

To investigate whether *Pf*MSP2 knock-out caused any changes in gene-expression of proteins that may have a role in merozoite invasion and possibly compensate for loss of *Pf*MSP2, we undertook RNA sequencing and differential gene expression analysis of *Pf*3D7 ΔMSP2 and *Pf*3D7 WT parasites. As expected, the *Pf*MSP2 transcript, which was targeted for removal by our CRISPR-Cas9 strategy, was not detected in the *Pf*3D7 ΔMSP2 parasites between 132 bp and 819 bp (Supplementary Figure 5). After removal of genes belonging to variant surface antigen families from the data, eight transcripts were found to have a log2fold expression increase above 1; none of these proteins are annotated as a merozoite surface or secreted protein linked to merozoite invasion (Figure 4C, Table 1). Five of these eight upregulated proteins are of unknown function and expressed at schizont stages, and only one (*Pf*3D7_0909100) has a transmembrane domain that could possibly anchor it to a membrane surface as the GPI anchor does for *Pf*MSP2 (Table 1). Three proteins were found to have a log2fold expression of <-1, indicating a significant reduction in expression with *Pf*MSP2 knock-out, with *Pf*MSP2 being one of these and the other two predicted to be exported proteins (Figure 4C, Table 1). While these data do not exclude any of the *Plasmodium* proteins with unknown function that are upregulated in *Pf*3D7 ΔMSP2 parasites as having a role that compensates for the loss of *Pf*MSP2, only one is predicted to have a transmembrane domain that would bind it to a membrane and no protein identified as up or down-regulated with *Pf*MSP2 knock-out has previously been linked to merozoite invasion. Therefore, *Pf*MSP2 knock-out does appear to have led to changes in gene expression, but does not appear to have a significant impact on the regulation of known merozoite surface or secreted protein gene-expression that would suggest loss of *Pf*MSP2 is compensated for by another protein.

### Impact of *Pf*MSP2 disruption on antibodies and inhibitors targeting secreted surface exposed antigens

Merozoites are key targets of antibodies during malaria infection with antibodies either able to directly block protein function and thus merozoite invasion, or recruit effectors such as complement leading to the destruction of the merozoite (Beeson et al., 2016). Given the prominence of *Pf*MSP2 on the merozoite surface we asked the question; how does the loss of *Pf*MSP2 affect antibody efficacy against merozoites? *Pf*EBAs and *Pf*Rhs moderate early steps in invasion with a degree of redundancy between these proteins. Several EBAs and Rhs can be targeted with invasion inhibitory antibodies and are targets of acquired human antibodies (Persson et al., 2013, 2008; Richards et al., 2013). We found no significant difference in the ability of purified rabbit immunoglobulin raised against single (*Pf*Rh2b) or combinations of *Pf*EBAs and *Pf*Rhs (*Pf*EBA175/*Pf*Rh2b, *Pf*EBA175/*Pf*Rh4, *Pf*EBA175/*Pf*Rh2b/*Pf*Rh4) to inhibit growth of *Pf*3D7 WT or *Pf*3D7 ΔMSP2 parasites (Figure 5A). Similarly, knock-out of MSP2 in *Pf*Dd2 did not significantly change the parasites sensitivity to rabbit antibodies raised against isogeneic W2mef EBA175 (Figure 5B). Broadly speaking, these results mirror what was seen with selective cleavage of the RBC ligands of *Pf*EBAs and *Pf*Rhs using enzyme treatment of RBCs (Figure 4B), with no evidence that loss of *Pf*MSP2 changed either the importance of these secreted merozoite antigens during invasion or their sensitivity to antibodies targeting different antigens.

**Figure 5.**
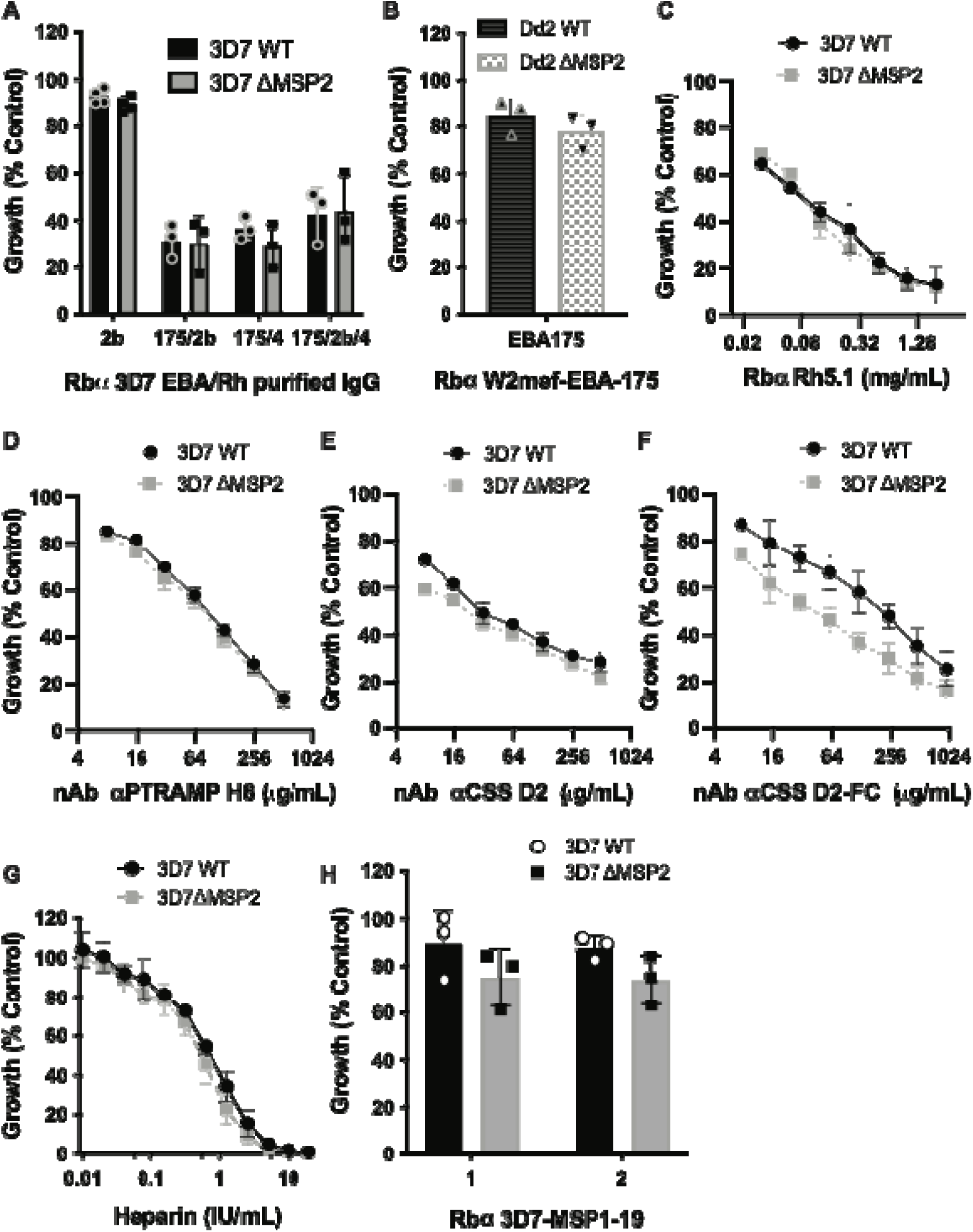
Impact of *Pf*MSP2 removal on efficacy of antibodies targeting other merozoite surface-exposed antigens. Changes in antibody efficacy in the absence of *Pf*MSP2 was assessed by measuring changes in antibody invasion inhibition and subsequent growth compared to growth in the absence of antibody for both *P. falciparum* 3D7 and Dd2 WT, and 3D7 ΔMSP2 Clone 1 and Dd2 ΔMSP2 Clone 1, parasites over 2 cycles. **(A)** Rabbit (Rb) IgG raised against merozoite antigens of the *Pf*3D7 EBA/Rh family. **(B)** Rabbit sera raised against *Pf*Dd2 EBA175. **(C)** Rabbit IgG raised against *Pf*3D7 Rh5. **(D)** Nanobody (nAb) to *Pf*3D7 PTRAMP. **(E-F)** Nanobody and Fc-tagged nanobody to *Pf*3D7 CSS. **(G)** The invasion inhibitory glycosaminoglycan heparin. **(H)** Rabbit sera raised against *Pf*3D7 MSP1-19 (different vaccinated rabbit sera identified by numbers). Graph displays mean ± S.D. of three different experiments. Significance was determined by unpaired t-test when only a single concentration point was tested and for IC_50_ comparisons an extra Sum-of-Squares F Test (best-fit LogIC_50_) was performed with p< 0.05 deemed significant.

The interaction of *Pf*Rh5 with RBC surface receptor basigin is a critical, non-redundant interaction for *P. falciparum* merozoite invasion and a known target of direct invasion-inhibitory antibodies (Alanine et al., 2019; Crosnier et al., 2011; Douglas et al., 2011). *Pf*Rh5 was the initial member of the PCRCR complex to be described, with subsequent studies demonstrating that antibodies against other members of this complex could also block merozoite invasion of the RBC (Scally et al., 2022). Given the importance of these secreted antigens to current vaccine development efforts, we next investigated whether loss of *Pf*MSP2 impacted on the parasite’s sensitivity to antibodies targeting these antigens. There was no difference in sensitivity between *Pf*3D7 WT and *Pf*3D7 ΔMSP2 parasites when treated with invasion-inhibitory rabbit antibodies targeting *Pf*Rh5 (Figure 5C) or nanobodies targeting PCRCR complex component *Pf*PTRAMP (H8) (Figure 5D). Nanobodies targeting the PCRCR complex component CSS (D2) showed a slight, but non-significant increase in invasion-inhibitory activity in the absence of *Pf*MSP2 (1.4-fold increase; IC_50_ value for *Pf*3D7 WT is 57 μg/mL; IC_50_ value for *Pf*3D7 ΔMSP2 is 42 μg/mL; p=0.3; Figure 5E). However, the *Pf*3D7 ΔMSP2 specific increase in invasion-inhibitory activity was amplified with the addition of the human Fc domain to the anti-CSS nanobody (D2-Fc) to be 3-fold greater for *Pf*3D7 ΔMSP2 parasites (IC_50_ *Pf*3D7 WT 138 μg/mL; IC_50_ *Pf*3D7 ΔMSP2 46 μg/mL; p=0.0001; Figure 5F).

The glycosaminoglycan heparin is a specific inhibitor of invasion and has been observed to bind to the 42 kDa C-terminal region of *Pf*MSP1 the most abundant GPI-anchored protein on the merozoite surface, marking this as a possible mechanism for its invasion inhibitory activity (Boyle et al., 2010a). However, the highly charged state of heparin means that it could block invasion through binding additional proteins. When tested against *Pf*3D7 WT and *Pf*3D7 ΔMSP2 parasites, we observed a small increase in inhibitory potency of heparin for parasites lacking *Pf*MSP2 (1.4-fold; IC_50_ *Pf*3D7 WT 0.65 IU/mL; IC_50_ *Pf*3D7 ΔMSP2 0.47 IU/mL; p=0.0003; Figure 5G). We next tested rabbit antibodies raised against the C-terminal 19 kDa fragment of *Pf*MSP1. We found a small but non-significant trend towards increased inhibition of growth in parasites that lacked *Pf*MSP2 across repeat experiments (Figure 5H). These data suggest that loss of *Pf*MSP2 can impact on the invasion-inhibitory potency of some antibodies that target secreted or surface antigens.

### Loss of *Pf*MSP2 consistently potentiates AMA1 invasion inhibitory antibodies

Subsequent to *Pf*Rh5 -basigin binding in the steps of merozoite invasion is the formation of the tight-junction between *Pf*AMA1, secreted from the micronemes onto the merozoite surface, and the rhoptry neck protein *Pf*RON2, which is inserted into the RBC membrane (Lamarque et al., 2011; Srinivasan et al., 2011). In lines where *Pf*MSP2 was disrupted, we found a consistent potentiation of antibodies that had anti-*Pf*AMA1 invasion-inhibitory activity. Invasion-inhibitory polyclonal rabbit serum antibodies and purified immunoglobulin raised against *Pf*AMA1 of *Pf*3D7 and *Pf*W2mef (isogenic to *Pf*Dd2), which express different alleles of both *Pf*MSP2 and *Pf*AMA1, were found to be more inhibitory with *Pf*MSP2 knock-out (Figure 6A and B). The polyclonal antibody with the greatest potentiation of inhibitory activity was an IgG preparation purified from a rabbit antiserum raised against *Pf*W2mef AMA1. This had a 3-fold increased potency against *Pf*Dd2 ΔMSP2 compared to *Pf*Dd2 WT (IC_50_ *Pf*Dd2 WT 0.015 mg/mL; IC_50_ *Pf*Dd2 ΔMSP2 0.005 mg/mL; p<0.0001) over two cycles of growth (Figure 6B, C).

**Figure 6.**
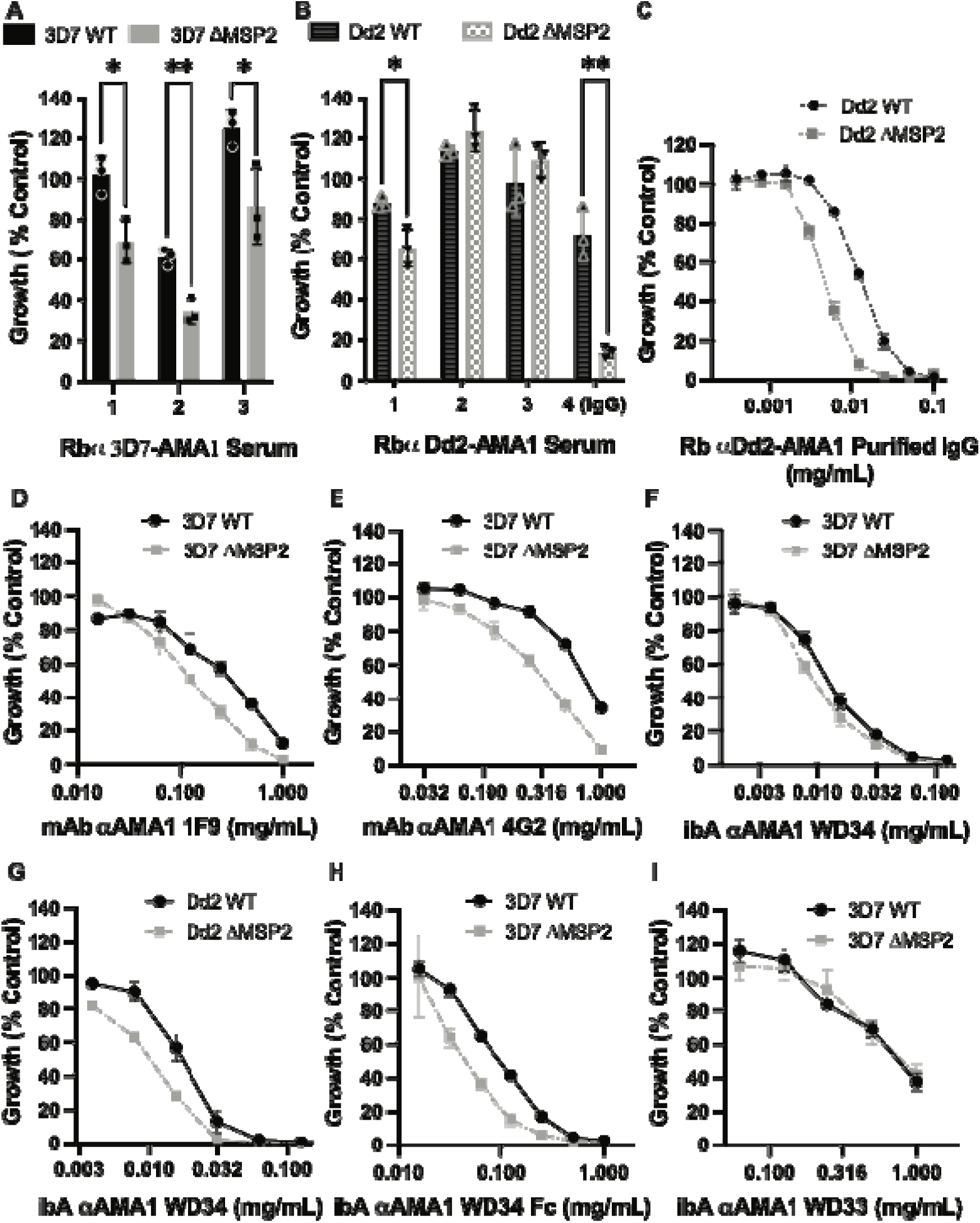
Absence of *Pf*MSP2 from the merozoite surface impacts invasion inhibition by *Pf*AMA1 antibodies. *Pf*3D7 **(A, D-F, H, I)** and *Pf*Dd2 **(B, C, G)** express different *Pf*MSP2 alleles and different *Pf*AMA1 alleles, yet both showed altered anti-AMA1 antibody growth inhibition for 3D7 ΔMSP2 Clone 1 and Dd2 ΔMSP2 Clone 1 parasites compared to parental parasites. The effect was seen with serum (**A-B**; different vaccinated rabbit (Rb) sera identified by numbers), purified rabbit and mouse monoclonal (mAb) antibodies **(C-E)** and i-bodies (ibA) **(F-J)**. Final parasitaemia was determined by flow cytometry and compared to control. Graph displays mean ± S.D. of three or four independent experiments. Significance was determined by unpaired t-test when only a single concentration point was tested and for IC_50_ comparisons an extra Sum-of-Squares F Test (best-fit LogIC_50_) was performed with p< 0.05 deemed significant.

The mouse 1F9 (binds to the RON2 binding pocket and polymorphic loop 1d (Coley et al., 2007)) and rat 4G2 (binds to a conserved epitope in the ectodomain (Kocken et al., 1998)) monoclonal antibodies have been shown to target different regions of *Pf*AMA1 and exhibit selective inhibition against *Pf*3D7 AMA1. Here we found potentiation of invasion inhibitory activity with *Pf*3D7 MSP2 knock-out for both 1F9 (2.1-fold; IC_50_ *Pf*3D7 WT 0.27 mg/mL; IC_50_ *Pf*3D7 ΔMSP2 0.13 mg/mL; p<0.0001; Figure 6D) and 4G2 (3.2-fold; IC_50_ *Pf*3D7 WT 1.15 mg/mL; IC_50_ *Pf*3D7 ΔMSP2 0.36 mg/mL; p<0.0001; Figure 6E).

We also tested the invasion inhibitory effect of i-bodies, which are smaller single-domain antibody-like molecules inspired from the shark variable new antigen receptor (V_NAR_) (Griffiths et al., 2018, 2016). When we tested the i-body WD34 (Angage et al., 2024) which binds a highly conserved epitope that includes the *Pf*RON2-binding pocket on *Pf*AMA1 domain II, we observed a small potentiation of *Pf*AMA1 specific activity with knock-out of *Pf*MSP2 in *Pf*3D7 (1.3-fold; IC_50_ *Pf*3D7 WT 0.012 mg/mL; IC_50_ *Pf*3D7 ΔMSP2 0.009 mg/mL; p=0.08 Figure 6F). Given WD34 was also found to be inhibitory to growth of W2mef parasites (Angage et al., 2024) we also tested its activity against our *Pf*Dd2 ΔMSP2 parasites and found potentiation of invasion inhibitory activity with loss of MSP2 in *Pf*Dd2 parasites (2-fold; IC_50_ *Pf*Dd2 WT 0.016 mg/mL; IC_50_ *Pf*Dd2 ΔMSP2 0.008 mg/mL; p=0.004 Figure 6G). As was observed for the anti-CSS nanobody, addition of a human Fc domain to WD34 amplified the inhibitory phenotype, with WD34 Fc having a 2.3-fold greater inhibitory activity against *Pf*3D7 ΔMSP2 than against *Pf*3D7 WT (IC_50_ *Pf*3D7 WT 0.1 mg/mL; IC_50_ *Pf*3D7 ΔMSP2 0.04 mg/mL; p=0.0004; Figure 6H). A second i-body, WD33 (Angage et al., 2024), which binds AMA1 between domain II and domain III but does not appear to overlap with the *Pf*RON2-binding pocket on *Pf*AMA1, had very limited invasion inhibitory activity against *Pf*3D7 parasites and did not show improved potency with knock-out of *Pf*3D7 MSP2 (0.9-fold; IC_50_ *Pf*3D7 WT 1.02 mg/mL; IC_50_ *Pf*3D7 ΔMSP2 1.1 mg/mL; p=0.8; Figure 6I). Although the limited inhibitory activity for WD33 tagged with the Fc receptor prevented us determining an IC_50_ at the concentrations feasible to test, there was no increased inhibition of *Pf*3D7 ΔMSP2 parasites compared to the low level seen with *Pf*3D7 WT parasites (data not shown). These data suggest that the potentiation of *Pf*AMA1 targeted antibody inhibition with MSP2 knock-out is *Pf*AMA1 epitope dependent.

To account for the differences in i-body sizes, we compared the WD34 and WD34 Fc i-bodies on a molar basis and found that they had similar activities against *Pf*3D7 MSP2 WT parasites (IC_50_ 0.94 μM and 1 μM respectively) indicating that the Fc-tag itself did not modify epitope binding properties. Rather, it is the absence of *Pf*MSP2 that results in increased inhibition of AMA1 function by WD34, and this is further potentiated by antibody size.

In light of these results, we hypothesised that removal of *Pf*MSP2 may improve antibody access to other antigens that are targets of inhibitory antibodies. To assess whether this was the case, we first tested whether increased *Pf*AMA1 polyclonal antibody binding to the surface of merozoites could be observed for *Pf*3D7 ΔMSP2 compared to *Pf*3D7 MSP2 WT parasites using a fluorescently tagged WD34 i-body (WD34-mCherry) for quantitative immunofluorescence assays of late stage schizonts labelled with a single incubation and wash step. We observed WD34-mCherry to have a significantly higher mean fluorescence intensity for *Pf*3D7 ΔMSP2 compared to *Pf*3D7 WT (Figure 7A), supporting that the *Pf*AMA1 i-bodies may have better access in the absence of *Pf*MSP2. As a control, we tested labelling using a fluorescently tagged WD33 i-body which binds to AMA1 (Angage et al., 2024), but we found did not have improved inhibitory activity in the absence of MSP2 (Figure 6I). Mirroring the results of the growth assay, there was no difference observed for WD33-eGFP binding fluorescence for *Pf*3D7 ΔMSP2 compared to *Pf*3D7 WT parasites (Figure 7B).

**Figure 7.**
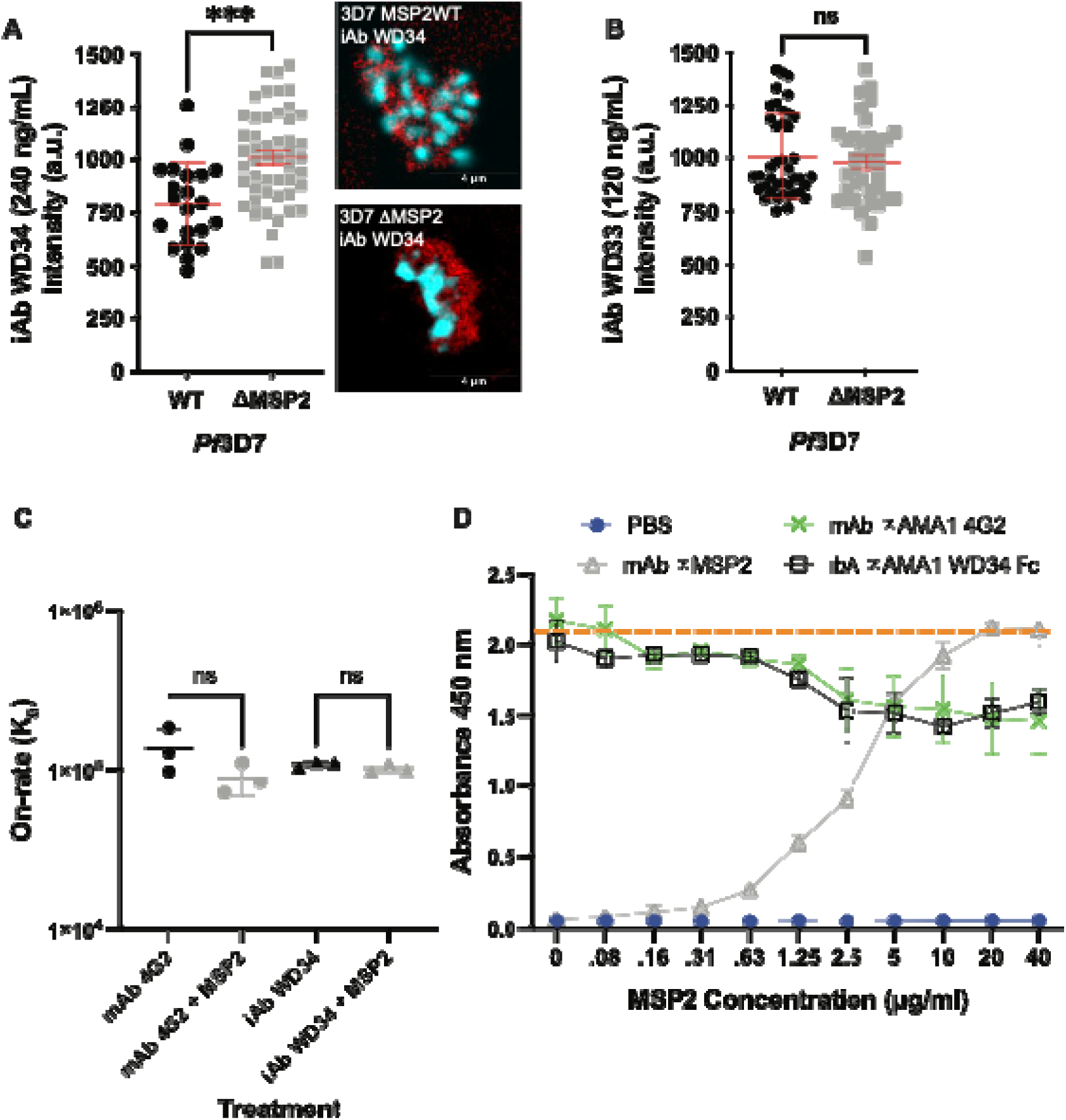
Quantitative fluorescence microscopy to assess whether differential binding may explain the increased potency of anti-*Pf*AMA1 invasion-inhibitory antibodies in the absence of *Pf*MSP2. Fluorescence intensity of fluorescently tagged anti-*Pf*AMA1 i-body (ibA) WD34-mCherry **(A)** and WD33-eGFP **(B)** for both *Pf*3D7 WT and *Pf*3D7 ΔMSP2 Clone 1 parasites, with a representative image for i-body WD34-mCherry. Nucleus in blue, i-body signal in red. Scale bar = 4 µm. Two independent experiments were performed with significance determined by unpaired t-test with p< 0.05 deemed significant. The lower overall mCherry signal required a higher antibody concentration (240 ng/mL) to have a comparable intensity measure to the eGFP tagged antibody (120 ng/mL) for *Pf*3D7 WT merozoites. **(C)** Read-out of the Surface plasmon resonance (SPR) antibody on-rate (association constant) for anti-*Pf*AMA1 mAb 4G2 and i-body WD34-Fc (mouse Fc) binding to *Pf*AMA1 in the presence or absence of *Pf*MSP2 protein. Data represents the mean of 3 experiments with significance determined by unpaired t-test with p< 0.05 deemed significant. **(D)** ELISA based assessment of the anti-*Pf*AMA1 mAb 4G2 and i-body WD34-Fc antibody binding levels to recombinant *Pf*3D7 AMA1 in the presence or absence of recombinant *Pf*3D7 MSP2. PBS control demonstrates background fluorescence. Dashed orange line provides a guide for peak absorbance levels. Anti-*Pf*MSP2 mAb shows increasing concentrations of *Pf*MSP2 protein results in decreased binding of mAb 4G2 and i-body WD34-Fc. Data represents the mean of 3 experiments and error bars are ±S.D.

We next explored whether the addition of recombinant *Pf*MSP2 could impact on the inhibitory antibody binding on-rate against substrate-bound *Pf*AMA1 recombinant protein using Surface Plasmon Resonance (SPR). For the anti-*Pf*AMA1 targeting mAb 4G2, there was a non-significant trend toward a lower on-rate in the presence of recombinant *Pf*MSP2 (Figure 7C). For the i-body WD34-Fc, there was minimal difference in detectable binding to *Pf*AMA1 in the presence or absence of *Pf*MSP2 (Figure 7C). Using a similar principle of testing antibody binding to substrate-bound *Pf*AMA1 protein in the presence, or absence of increasing concentrations of *Pf*MSP2, we assessed changes in levels of anti-*Pf*AMA1 antibody binding using an enzyme linked immunosorbent assay (ELISA). With increasing concentrations of *Pf*MSP2 added to wells coated with AMA1, we found decreased binding of anti-*Pf*AMA1 mAb 4G2 and i-body WD34-Fc (mouse Fc) by ELISA (Figure 7D). Concentrations where the *Pf*MSP2 protein was barely detectable above background using an MSP2 mAb were found to be enough to impact on the binding of *Pf*AMA1 antibodies. In contrast, increasing concentrations of the intrinsically disordered MSP4 from *P. falciparum* 3D7 (40 kDa) and the highly structured immunoglobulin domain of neural cell adhesion molecule (NCAM, CD56, 16 kDa) recombinant proteins did not impact on binding of anti-*Pf*AMA1 antibodies to recombinant AMA1 (Supplementary Figure 6). By looking at the impacts of removing *Pf*MSP2 from the merozoite surface, or adding this protein in to recombinant assays, these data provide additional support that inhibitory antibodies targeting *Pf*AMA1 bind at a higher rate in the absence of *Pf*MSP2.

## Discussion

Despite decades of research and interest, the function of most merozoite surface proteins remains unclear and targeting merozoite antigens in vaccine development has not achieved high efficacy in clinical trials. Additionally, it has generally proven difficult to generate highly inhibitory antibodies through vaccination with merozoite antigens. *Pf*MSP2 is one of the most abundant proteins on the merozoite surface, has shown promise as a vaccine candidate and is an established target of protective cytophilic antibodies (Boyle et al., 2015; Genton et al., 2002; McCarthy et al., 2011; Osier et al., 2014, 2010). However, its function remains unclear. Our findings here suggest that *Pf*MSP2 does not have an essential role in invasion *in vitro*, but can modulate the sensitivity of merozoites to inhibitory antibodies targeting other antigens.

Despite the abundance of *Pf*MSP2 on the merozoite surface and previous work suggesting a role in RBC invasion, we found merozoites invade and grow with similar kinetics to wildtype parasites in the absence of *Pf*MSP2. This does not exclude a role for *Pf*MSP2 *in vivo* where there are additional pressures, such as immune-effector mechanisms and flow dynamics, on merozoite invasion. However, given we have knocked-out *Pf*MSP2 from two different *P. falciparum* isolates, our findings do not currently support a major role for *Pf*MSP2 in the mechanics of merozoite invasion. Thus, it appears that the function of the two most abundant proteins on the merozoite surface, *Pf*MSP1 (Das et al., 2015; Kals et al., 2024) and *Pf*MSP2, are not obviously linked to merozoite binding to the RBC and subsequent invasion. Deletion of MSP2 did lead to significant changes in the expression of several genes, but there were no proteins with a clear link to merozoite invasion and the biological relevance of these changes is not currently understood and requires future investigation.

Both rabbit polyclonal and mouse monoclonal anti-*Pf*AMA1 antibodies tested were consistently more inhibitory in the absence of *Pf*MSP2. Of the two *Pf*AMA1 targeting i-bodies tested (Angage et al., 2024), the most potent WD34 was also more potent in the absence of *Pf*MSP2, with this activity increasing significantly with the ∼5.6-fold increase in size that occurred upon addition of a human Fc region. Similarly, activity of a nanobody targeting the PCRCR complex protein CSS showed increased potency in the absence of *Pf*MSP2, and again this was increased further with antibody enlargement through addition of an Fc region. These data suggest that *Pf*MSP2 may act to dampen the inhibitory activity of antibodies targeting other antigens on the surface of the merozoite. Such immune dampening (called conformational masking) by intrinsically disordered proteins, or protein regions, of host-cell invasion related proteins has recently been demonstrated by modelling and gene-editing studies in the viral pathogens HIV and HCV, respectively (Li et al., 2024; Stejskal et al., 2022). These studies, along with our demonstration of increase antibody inhibitory efficacy with loss of the intrinsically disordered *Pf*MSP2, support that this may be a widespread approach used by pathogens to protect important protein functional sites from inhibitory antibodies.

A role in immune evasion has previously been suggested for *Pf*MSP2 and speculatively an alternate explanation for the sensitisation observed is that this protein may shield key *Pf*AMA1 epitopes from antibody inhibition. For the *P. falciparum* 3D7 line, we found that *Pf*MSP2 knock-out led to higher *Pf*AMA1 binding as measured by increased fluorescence intensity for the mCherry tagged i-body WD34 in the absence of *Pf*MSP2. Using a solid-phase model and titrating recombinant *Pf*MSP2, we found that increasing concentrations of *Pf*MSP2 reduced binding of an anti-*Pf*AMA1 mAb and i-body, even at concentrations where *Pf*MSP2 was not detectable itself by antibodies. These data support that the presence of *Pf*MSP2 may reduce antibody access (masking) to important invasion proteins exposed on the merozoites surface. However, we acknowledge that the antibody-binding assays used (immunofluorescence assays, SPR, ELISA) might not be sensitive enough to observe all potential changes in binding, temporal changes when binding epitopes are exposed or differences in avidity that could lead to increased antibody potency during the course of parasite invasion and growth, and these factors may also contribute. Future studies of the merozoite surface and interactions between proteins may help decipher in full how *Pf*MSP2 is protecting essential merozoite antigens from invasion inhibitory antibody activity and reveal strategies for enhanced vaccine design.

Our observation that loss of *Pf*MSP2 potentiates the invasion-inhibitory activity of antibodies targeting other antigens opens an additional avenue for vaccine development, such as targeting *Pf*MSP2 to interfere with its potential role in immune evasion and thus potentiating antibodies against other merozoite antigens. Additionally, our results suggest that vaccines based on *Pf*AMA1 may need to be designed to specifically avoid the effect of *Pf*MSP2 modulating inhibitory antibodies, such as focussing on specific epitopes or structural features. To date, vaccines based on AMA1 have failed to achieve substantial efficacy (Sagara et al., 2009; Sheehy et al., 2012; Thera et al., 2011). We can speculate that one possible protective mechanism of action of the Combination B vaccine (Genton et al., 2002; McCarthy et al., 2011), which combined *Pf*MSP2, *Pf*MSP1 and *Pf*RESA, could be that the inhibitory activity of vaccine-induced antibodies raised against *Pf*MSP2 were then able to potentiate the activity of antibodies targeting other antigens, potentially *Pf*MSP1 which was also a target of the vaccine. Here we show consistent potency improvement with *Pf*MSP2 knock-out for growth inhibitory rabbit, mouse monoclonal and i-body antibodies targeting *Pf*AMA1, as well as demonstrate improved activity for and Fc-tagged nanobody targeting *Pf*CSS, indicating that these are not outlier results from a single antibody or antibody type. However, increased antibody potency was not shared across all antibodies tested, possibly because the specific function or localisation of a target protein, the region that an antibody binds to or the functional activity (or lack thereof) of an antibody may all play a role in determining whether loss of *Pf*MSP2 can potentiate growth inhibitory activity. Further investigation using the parasite lines developed in this study and a wider panel of antibodies that target different stages of the merozoite invasion process, including human monoclonal antibodies against AMA1(Patel et al., 2025), could shed more light on this potentially novel mechanism of vaccine derived antibody efficacy.

The results of this study and work done by Escalante et al. (2022) demonstrate that MSP2-like sequences are found in the avian malarias *P. galinaceum* and *P. relictum*, which differs from the fields previous understanding that *Pf*MSP2 arose in the *Laverania* lineage of malaria parasites. This observation indicates that MSP2 was likely present in the ancestral malaria parasite before being lost in primate and rodent *Plasmodium* spp. Further insights into the evolution and loss of *Pf*MSP2 are likely to be found with additional whole genome sequences of bird and lizard malaria species. The presence of *msp2*-like sequences in some avian *Plasmodium* species and *Laverania* suggest a retention of *msp2* sequences in the *Laverania* which indicates an important role for MSP2 for these parasites. Indeed, our amino acid and structural prediction level comparison demonstrates that much of the protein properties conserved in the *Laverania* MSP2s are also recognisable, and in many cases conserved, in the avian malaria MSP2s, despite 10 million years of evolutionary divergence. This suggests that having an MSP2-like protein is beneficial for these parasites with very different hosts. However, CRISPR-Cas9 gene editing used in this work has shown that, in contrast to previous attempts to knock-out *Pf*MSP2 (Sanders et al., 2006), *Pf*MSP2 is not essential for *P. falciparum* blood stage parasite growth *in vitro*. Our findings raise the possibility that the retention of *Pf*MSP2 in the *Laverania*, and by association potentially the avian malaria parasites, may be linked to its apparent capacity to modulate the impact of antibodies against other merozoite surface-exposed antigens, rather than through a direct mechanistic role in host-cell invasion. If MSP2 was to fulfill such a role in the *Laverania* and avian malaria parasites, it raises the question of whether a similar protective system exists in other *Plasmodium* spp. and what the key protein/s are that fulfil this role in these parasites.

The *Pf*3D7 MSP2 and *Pf*Dd2 MSP2 lines developed in this study will provide useful reagents for investigating the potential of targeting a single, or both, *Pf*MSP2 allelic types with vaccines that elicit different immune responses. These parasites could also be used to explore whether vaccine-induced antibodies against other merozoite antigens could also be potentiated if any immune evasion mechanism provided by *Pf*MSP2 is removed. This may be achievable with a combination vaccine or through targeting an epitope of another merozoite vaccine candidate that is not protected by *Pf*MSP2. Such information, investigated using the *Pf*MSP2 knock-out parasites developed in this study, could be useful in prioritising vaccine candidates, specific epitopes and combinations that target the merozoite.

## Conclusion

Advancements in gene-editing techniques in *P. falciparum* have allowed us to directly demonstrate using reverse genetics in two different parasite lines that *Pf*MSP2 is not essential for *P. falciparum* growth *in vitro*. Instead, we present a new concept that MSP2 can modulate the activity of invasion-inhibitory antibodies and may have evolved for this purpose. It is possible that other merozoite surface proteins may have similar roles and should be investigated in future studies. Our observation that loss of *Pf*MSP2 potentiates the inhibitory activity of antibodies targeting other merozoite surface exposed proteins reveals a new avenue to explore in vaccine development by blocking or bypassing this function to improve the activity of vaccines targeting these antigens.

## Materials and Methods

### Bioinformatic analysis of *Pf*MSP2, *Pf*MSP4 and *Pf*MSP5 like proteins in *Plasmodium* spp

The PlasmoDB database (Aurrecoechea et al., 2009) was used to identify in the available *Plasmodium* genomes the region between Adenylosuccinate lyase (PF3D7_0206700) and a conserved protein of unknown function (PF3D7_0207100) where *Pf*MSP2, *Pf*MSP4 and *Pf*MSP5 are localised in *P. falciparum*. All annotated MSP2, MSP4 and MSP5 protein sequences were downloaded. Translated sequences of unannotated genes and putative annotations in this region were also downloaded and aligned using the MUSCLE alignment tool MEGA11: Molecular Evolutionary Genetics Analysis version 11 (Tamura et al., 2021) or the Geneious Prime 2019.2.1 MUSCLE alignment tool (Biomatters Ltd.) and determined to be MSP2, MSP4 or MSP5 based on sequence similarity in the conserved regions of these genes. A BLAST (NCBI) search with the N-terminal and C-terminal conserved regions of MSP2 was also performed to capture any sequences missed by the above approach. A maximum likelihood phylogenetic tree of identified MSP2 sequences was generated using the default maximum likelihood settings of MEGA11 with 500 bootstraps and the *P. gallinaceum* MSP2 as an outgroup.

### Modelling of MSP structure across *Plasmodium* spp. using Alpha-fold

In order to obtain models of the 3-dimensional protein structures of MSP2 for *P. falciparum*, *P. reichenowi*, *P. billcollinsi*, *P. adleri*, *P. relictum* and *P. gallinaceum*, we first removed the predicted N-terminal signal sequences and C-terminal GPI anchor sequence. The resulting sequences were submitted to AlphaFold2 (Jumper et al., 2021). The resulting models were subsequently analysed and visualised using PyMOL 2.4.0 (Schrödinger, USA).

### Culture and synchronisation of *P. falciparum*

*Plasmodium falciparum* 3D7 and Dd2 was cultured in RPMI-HEPES media (Gibco) containing 0.5% Albumax ll (Gibco), 25 μM Gentamicin (Gibco), 367 μM hypoxanthine (Sigma Aldrich), 2 mM L-Glutamax (Gibco), 0.17% sodium bicarbonate (Sigma Aldrich) and O positive human red blood cells (Lifeblood, Australia) at 3% parasitemia, 3% haematocrit. Cultures were grown at 37°C under 1% oxygen, 5% CO_2_ (BOC gas), in a humidified chamber. For gene-edited lines, WR99210 (Jakobus Pharmaceuticals) was added to culture media. Sorbitol lysis (5%) (Sigma Aldrich) to select for ring-stage parasites, Percoll Plus (Sigma Aldrich) selection of late-stage parasites and heparin treatment (Pfizer) (Boyle et al., 2010b) were used to maintain synchronicity of parasite cultures.

### Generation of *Pf*MSP2 knockout line using CRISPR Cas9

Briefly, a flank region incorporating the 5’ region upstream of *Pf*MSP2, the first 44 amino acids of the protein, and a flank of the region immediately 3’ of *Pf*MSP2 was inserted into the pCC1 plasmid by restriction enzyme cloning. A 20 bp guide sequence was designed using EuPaGDT (Peng and Tarleton, 2015), annealed and inserted in the pUF_Cas9 plasmid by infusion cloning (Takara Bio). pUF_Cas9 guide plasmid (20 μg) and pCC1 *Pf*MSP2 repair plasmid (60 μg) was transfected directly into *P. falciparum* schizonts. Early schizonts were purified using a 70% percoll gradient and treated for 4 hrs with compound 1 (Taylor et al., 2010) at 2 μM. After incubation treated schizonts were washed twice and left shaking for 20 minutes before pelleting and resuspension in Complete Cytomix (plasmids plus 0.895% KCl, 0.0017% CaCl_2_, 0.076% EGTA, 0.102% MgCl_2_, 0.0871% K_2_HPO_4_, 0.068% KH_2_PO_4_, 0.708% HEPES). Schizonts were electroporated in 0.2 cm cuvettes (Biorad) at 800 V and 25 μF. Electroporated schizonts were mixed with warmed media and fresh RBCs and shaken to promote invasion for 20 minutes after which they were placed in a culture dish. Transfected parasites were then treated with WR99210 to select for integration of the hDHFR drug selection cassette and MSP2 knock-out.

To confirm integration gDNA was collected by saponin lysis of schizont stage parasite culture and gDNA extracted using the PureLink Genomic DNA Mini Kit (Invitrogen) as per manufacturer’s instructions. Integration of the *Pf*MSP2 knockout construct into a portion of parasites was confirmed by PCR (Supplementary Table 1). Once confirmed cultures were cycled on and off WR99210 (Jakobus Pharmaceuticals) and 1 μM 5FC (Sigma Aldrich) to select for integrated parasites no longer expressing the guide plasmid. Limited dilution cloning was then performed to obtain parasite clones which had integrated the knockout construct.

### Western blotting to detect *Pf*MSP2

Synchronised schizonts were harvested and the RBCs lysed by saponin. Briefly ∼10 mL schizont (38-44 hrs) culture was lysed on ice for 10 minutes with 0.15% w/v saponin before pelleting by centrifugation and washing once in 0.075% w/v saponin followed by three washes in PBS containing protease inhibitors (CØmplete, Roche). Saponin treated schizont pellets were treated with DNAsel (Qiagen) for 10 minutes at room temperature before resuspension in reducing sample buffer (0.125 M Tris-HCL, 4% SDS, 20% glycerol, 10% beta-mercaptoethanol and 0.002% bromophenol blue). Proteins were separated by size using SDS-PAGE 4-12% Bis-Tris Gels (Bolt, Invitrogen) at 110V for 80 minutes before transfer to a nitrocellulose membrane (iBlot, Invitrogen) at 20V for 7 minutes. Blots were blocked from 1 hr to overnight in 3% skim milk PBS (Sigma Aldrich) before incubation with primary antibodies (1/75000 mouse 2F2 anti-*Pf*MSP2 3D7 (Supplementary Table 2; (Adda et al., 2012)) and 1/75000 rabbit anti-*Plasmodium* aldolase (Abcam) or 1/75000 rabbit anti-FC27 and 1/10000 mouse anti- EXP2 (a gift of Paul Gilson, Burnet Institute, Melbourne) for 1- 2 hrs. IRDye 800CW goat anti-mouse (1/4000, LI-COR Biosciences) or IRDye 680RD goat anti- rabbit (1/4000, LI-COR Biosciences) secondary antibodies were used for detection on the Odyssey Infrared imaging system (LI-COR Biosciences). Quantification was performed using Image Studio Lite 5.2.5 (LI-COR Biosciences).

### Quantification of parasite expansion rate and invasion inhibition

Potential growth defects resulting from gene editing was assessed by flow cytometry. The initial parasitemia of cultures was determined by flow cytometry and then measured again after the 50 μL cultures in 96 well plates were maintained under standard (still) or shaking (50 rpm) conditions for 48 hrs or 96 hrs of growth. To determine parasitemia by flow cytometry Ethidium Bromide (10 μg/mL BioRad) was added to 50 μL of 1% haematocrit culture for 30 mins to allow for staining of parasite DNA. The stain was washed off and wells resuspended in 200μL of PBS. Data was collected on a BD Accuri C6 Plus Flow Cytometer (BD Biosciences) and analysed on FlowJo (FlowJo LLC). Briefly, forward scatter and side scatter was used to determine the RBC population. From this population infected RBCs were gated on as highly fluorescent in the PE channel. Final parasitemia was compared back to initial parasitemia to determine the fold increase in parasitemia for each line.

To examine the invasion inhibitory effect of antibodies (Supplementary Table 2), 5 μL of potential inhibitor was mixed with 45 μL of 0.2% schizont parasitemia at 1% haematocrit for 96 hrs before end parasitemia was determined by flow cytometry. Data is displayed as % Growth of media only controls for each line.

To examine differences in RBC receptor preference RBCs were treated with different enzymes that cleave different residues of RBC receptors which is used as a marker for which RBC receptor/s are preferred by *P. falciparum* (Duraisingh et al., 2003). Infected RBCs at 1% ring stage parasitemia were washed three times in RPMI-HEPES to remove excess protein. Packed RBCs (20 μL) at 1% parasitemia were then resuspended with 5X RBC volume of pre-warmed enzymes at desired concentration (Supplementary Table 3). RBC-enzyme mix was incubated at 37°C for 45 mins with rotation to keep RBCs resuspended. Following incubation, RBCs were washed three times and added to fresh complete media at 1% haematocrit in a U- bottom 96 well plate in technical duplicate. Plates were incubated for 72 hrs, sufficient time for *P. falciparum* to complete one full cycle of growth and reach trophozoite stage again. Final parasitemia was determined by flow cytometry, as detailed above, and compared to non-treated controls. IC_50_ calculations data was log transformed then a nonliner regression log-(inhibitor)-versus-response curve was calculated with Extra Sum-of-Squares F Test (best-fit LogIC_50_) used to compared IC_50_s between conditions.

### Immunofluorescence assay of Wildtype and *Pf*MSP2 KO parasites

Schizonts ∼38 hrs old were treated with E64 (Sigma-Aldrich) for 5 hrs to allow development of very mature schizonts desired for imaging. After E64 treatment, parasite cultures were fixed in 4% v/v paraformaledehyde (PFA, Sigma-Aldrich), 0.0075% v/v glutaraldehyde (pH 7.5, Electron Microscopy Sciences) solution for 30 min at room temperature shaking gently. Fixed parasites were washed in 1X PBS and then resuspended in PBS at 1% haematocrit. Coverslips (#1.5H high-precision coverslips, Carl Zeiss, Oberkochen, Germany) were coated in 0.01% Poly-L-lysine (Sigma Aldrich) for 1 hr at room temperature, washed in miliQ water before the fixed parasite culture was allowed to adhere for 1 hr. Cells were permeabilised by 0.1% Triton X-100 PBS for 10 mins before blocking for 1 hr to overnight in 3% BSA 0.05% Tween 20 PBS. Primary antibodies were diluted in 1% BSA 0.05% Tween 20 PBS and incubated for 2 hrs. Coverslips were washed in 0.05% Tween 20 PBS three times before incubation with the secondary antibody (1/500 goat anti-chicken/mouse/rabbit Alexa Fluor coupled secondary antibodies -488 nm, 594 nm, 647 nm; Life Technologies) for 1 hr. After the secondary antibody was washed off coverslips underwent a secondary fix in 4% v/v PFA, 0.0075% v/v glutaraldehyde for 5 mins. The fixative was washed off and the coverslips dehydrated with sequential 3 min ethanol treatment (70%, 90% and 100%). Coverslips were allowed to dry and mounted with ProLong Gold antifade solution (refractive index 1.4) which contains 4’, 6-diamidino-2phenylindole, dihydrochloride (DAPI) (ThermoFisher Scientific) and allowed to set overnight. Imaging was performed on the Zeiss LSM 800 using the Airyscan super-resolution mode (Carl Zeiss, Obekochen, Germany).

### Live cell microscopic analysis of merozoite RBC invasion

Highly synchronised *Pf*Dd2 and *Pf*Dd2 ΔMSP2 parasites at schizont stages were adjusted to 0.25% Haematocrit in complete culture medium. A volume of 200 μL was added to a well of an iBidi 15 μ-Slide 8 well, glass bottom slide (iBidi 80827) and the slide was promptly returned to sit on a prewarmed water block within a gassed box and left to incubate at 37°C to allow iRBCs to settle. The slide was then transported directly to a prewarmed Nikon TiE microscope with an environmental chamber heated to 37^ο^C and gas mixture of 1% O_2_, 5% CO_2_ and 94% nitrogen. Filming was conducted using either a 60X water or 100X oil objective. Differential interference contrast (DIC) imaging was captured at 3 – 3.5 Volts with a camera exposure of 60 – 180 milliseconds.

Captured footage was processed using the Nikon analysis software (NIS-Elements AR Analysis version 5.21.01) and analysis of the schizont rupture and merozoite invasion events was performed manually. The merozoites which invaded successfully were tracked and attachment time, reorientation time (if obvious), deformation start and end times, the deformation score (Weiss et al., 2015; Wilson et al., 2015), invasion initiation and completion times and echinocytosis start times were recorded.

### RNA extraction

Highly synchronised cultures were pelleted, resuspended in TriZol (Invitrogen) and incubated for 5 minutes at 37°C. Chloroform (Sigma Aldrich) was added, mixed and then spun at 12,000g for 30 minutes at 4°C. Supernatant was collected and mixed with an equal volume of 70% Ethanol and processed using the RNeasy Kit (Qiagen) to extract purified RNA. RNA was DNAse l (Qiagen) treated for 30 mins at room temperature and cleaned up on RNeasy columns according to the manufacturers protocol. Absence of gDNA was checked by qPCR of gDNA with primers (*Pf*SUB1) and DNAse I treatment repeated if necessary.

### qPCR of gene expression in schizonts

DNAse treated RNA was added to 40 mM dNTPs (Qiagen), 0.4 mg/mL random hexamer (Qiagen) and incubated at 65°C for 5 mins. After incubation 5x Superscript Buffer (Invitrogen), 100 mM DTT (Invitrogen), 40 U/μL RNAseOUT (Invitrogen) and 200 μ/mL Superscript lll Reverse Transcriptase (Invitrogen) was added and cycled 25°C 5 mins, 50°C for 60 mins and 70°C for 15 mins. Primers (Supplementary Table 4) were designed to detect *Pf*MSP2, *Pf*MSP4, *Pf*MSP5, *Pf*SUB1 and Fructose Bisphosphate Aldolase as a reference gene (Salanti et al., 2003). qPCR was performed on cDNA from WT and knockout parasites with PowerUp SYBR Green Master Mix (Applied Biosystems) and cycling parameters; 95°C for 10 mins followed by 40 cycles 95°C for 15 mins, 60°C for 14 mins before a dissociation step at 95°C for 2 mins followed by 60°C for 2 mins with a 2% ramp to 95°C for 2 mins on the QuantStudio 7 Flex System (ThermoFisher). qPCR was performed in triplicate for each of the three independent experiments. Change in gene expression in knockout lines compared to WT was quantified by the 2^-ΔΔCt^ method (Livak and Schmittgen, 2001).

### Differential gene expression in *Pf*MSP2 KO compared to WT schizonts

RNA sequencing of the schizont stage transcriptome of *Pf*3D7 MSP2 KO and *Pf*3D7 WT was performed using the Illumina total RNA with Ribo Zero plus for library preparation and sequenced on the NovaSeq 6000 platform (Victorian Clinical Genetics Services) with 2x150 bp paired end reads (Supplementary Table 5). Reads were aligned to the *Pf*3D7 reference genome (PlasmoDB v61) using STAR aligner (Dobin et al., 2013). Reads were called using Rsubread FeatureCounts (Liao et al., 2014) with reads counted by CDS and summarised by gene. Differential gene analysis was performed with DESeq2 (Love et al., 2014) and visualized in R studio using ggplots2 (Wickham, 2016). MSP2 RNASeq data is available at ArrayExpress through accession number E-MTAB-15427.

### Quantitative Immunofluorescence assay

Synchronous *Pf*3D7 MSP2 KO and *Pf*3D7 WT parasite cultures were treated with ML10 (Ressurreição et al., 2020) to allow mature schizonts to develop. Thin blood smears were then made and fixed in 100%, -20°C methanol for 5 mins. Blocking of smears was performed in 1% BSA PBS, then fluorophore conjugated WD34 (240 ng/mL) and WD33 (120 ng/mL) i-bodies (Angage et al., 2024) in 1% BSA PBS were added and incubated at room temperature for 1 hr. Smears were then washed, ethanol dehydrated and a coverslip mounted (ProLong Diamond Antifade mountant with DAPI (refractive index 1.47, ThermoFisher Scientific)) and allowed to cure for 48 hrs at room temperature. Fluorescence images were acquired using an Olympus FV3000 confocal microscope equipped with a 100x oil objective (NA 1.4). Quantification of parasite associated fluorescence was performed using a modified image analysis pipeline (Liffner et al., 2020) implemented in Imaris^TM^ (9.9 version) where non-specific background fluorescence was removed using rolling ball background subtraction and a thresholding algorithm applied to the image to generate binary masks, separating objects exceeding the threshold (putative parasites) from the background. Binary masks were further refined to exclude objects smaller or larger than a defined size range (< 3 μm and > 15 μm), corresponding to the expected size of AMA i-body stained schizonts. Following segmentation the surface area containing antibody signal were calculated and these values compared between *Pf*3D7 MSP2 KO and *Pf*3D7 WT parasites.

### Surface Plasmon Resonance (SPR) Assay

Surface plasmon resonance (SPR) was employed to determine the association rate constant (K_a_) of AMA1-antibody interactions in the presence of MSP2 using the Biacore T200 system (Cytiva). Flow cells 1, 2, and 3 were activated for 14 minutes using a 1:1 mixture of 0.1 M N-hydroxysuccinimide (NHS) and 0.4 M N-(3-dimethylaminopropyl)-N’-ethylcarbodiimide hydrochloride (EDC) at a flow rate of 5 μL/min. Flow cell one was immobilised with bovine serum albumin (BSA) to serve as a reference surface. AMA1 was immobilised at a concentration of 80 μg/mL in 10 mM sodium acetate (pH 4.5) to achieve a surface density of 1000 response units (RU) on flow cells Two and three. MSP2 was co-immobilised in flow cell three to a surface density of 2000 RU. All surfaces were subsequently blocked with a 7-minute injection of 1 M ethanolamine (pH 8.0). Purified antibodies were prepared in PBS containing 0.005% Tween-20 (pH 7.4) and injected over the flow cells at a flow rate of 60 μL/min at 25°C. Single-cycle kinetics, using a bivalent analyte model, was applied to analyse the interactions.

### Enzyme-Linked Immunosorbent Assay (ELISA)

ELISAs to evaluate the effect of MSP2 on the binding of AMA1-specific antibodies were carried out in a volume of 100 μL per well, either at room temperature for 1 hr or overnight at 4°C. All washing steps consisted of three washes with PBS containing 0.1% Tween-20 (PBS-T). Nunc Maxisorp plates (Thermo Fisher Scientific, USA) were coated with AMA1 at a concentration of 0.8 μg/mL, followed by washing and subsequent coating with increasing concentrations of recombinant HIS-tagged *Pf*MSP2, HIS-tagged *Pf*MSP4 or HIS-tagged NCAM. Wells were then blocked with 5% (w/v) skim milk in PBS and then incubated for 1 hr at room temperature with one of mouse Fc-conjugated WD34 anti-AMA1 i-body (Angage et al., 2024), mouse monoclonal anti-AMA1 antibody (mAb) 4G2 (Kocken et al., 1998), anti-MSP2 mAb 9G8 (Adda et al., 2012), anti-HIS antibody (Sigma Aldrich) to detect *Pf*MSP4 or anti-NCAM antibody (21H5, (Griffiths et al., 2016)) at a concentration of 2.5 μg/mL. The primary antibody was washed off and the HRP-conjugated secondary antibody (1:5000, Sigma Aldrich) was then applied for 1 hr at room temperature before washing. For signal development to detect bound primary antibodies, wells were incubated with 1-Step^TM^ Ultra TMB-ELISA Substrate Solution (Thermo Fisher Scientific, USA) according to the manufacturers instructions. The reaction was stopped by adding 1 M sulfuric acid, and the absorbance at 450 nm was measured using a microplate reader.

### Statistical Analysis

Three independent experiments were performed for all experiments in technical duplicate, unless otherwise stated. Data was graphed and statistical tests performed in Prism GraphPad v9 or R.

## Acknowledgements

The authors would like to thank Prof Alan Cowman, Dr Stephen Scally and Jennifer Thompson (WEHI, Melbourne) for provision of PCRCR complex antibodies. Assoc Prof Paul Gilson (Burnet Institute, Melb) for EXP2 and MSP1 antibodies. Dr Chris MacRaild (Monash University) for helpful discussion. Dr Adam Thomas for the *Pf*MSP4 protein (Burnet Institute). Dr Sonja Frölich and Dr Ben Liffner (Adelaide University) for assistance on quantitative immunofluorescence assays and Adelaide Microscopy for training and use of microscopy instruments. We thank Lifeblood, Australia, for provision of RBCs. DWW was supported by a Hospital Research Foundation (C-MCF-52-2019) and Australian Research Council (FT240100420) Future Fellowship. DWW and JGB by a Hospital Research Foundation Collaborative Grant (S-03-EOI-2021). JGB was supported by the National Health and Medical Research Council of Australia (Research Fellowship and Investigator Grant to JGB, Australian Centre for Research Excellence in Malaria Research). IGH and JC were supported by an Australian Research Council RTP scholarship. The Burnet Institute is supported by the NHMRC for Independent Research Institutes Infrastructure Support Scheme and the Victorian State Government Operational Infrastructure Support.

## Author Contributions

Conceived and designed the experiments: DWW, IGH, JC, JGB, DA, MF, RA. Performed the experiments and phylogenetic analysis: IGH, JC, KHL, OR, DA, KRT, NB. Analysed the data: IGH, DWW, JC, KHL, OR, DA, MF. Provided key discussions on PfMSP2 and/or key reagents: RA, MF, DA. Paper writing: DWW, JC, IGH, JGB with critical input from RA, MF, DA, OR, KRT and based on the PhD thesis of IGH.

