## Supplementary Figures for "An abundant merozoite surface protein of *Plasmodium falciparum* modulates susceptibility to inhibitory antibodies"

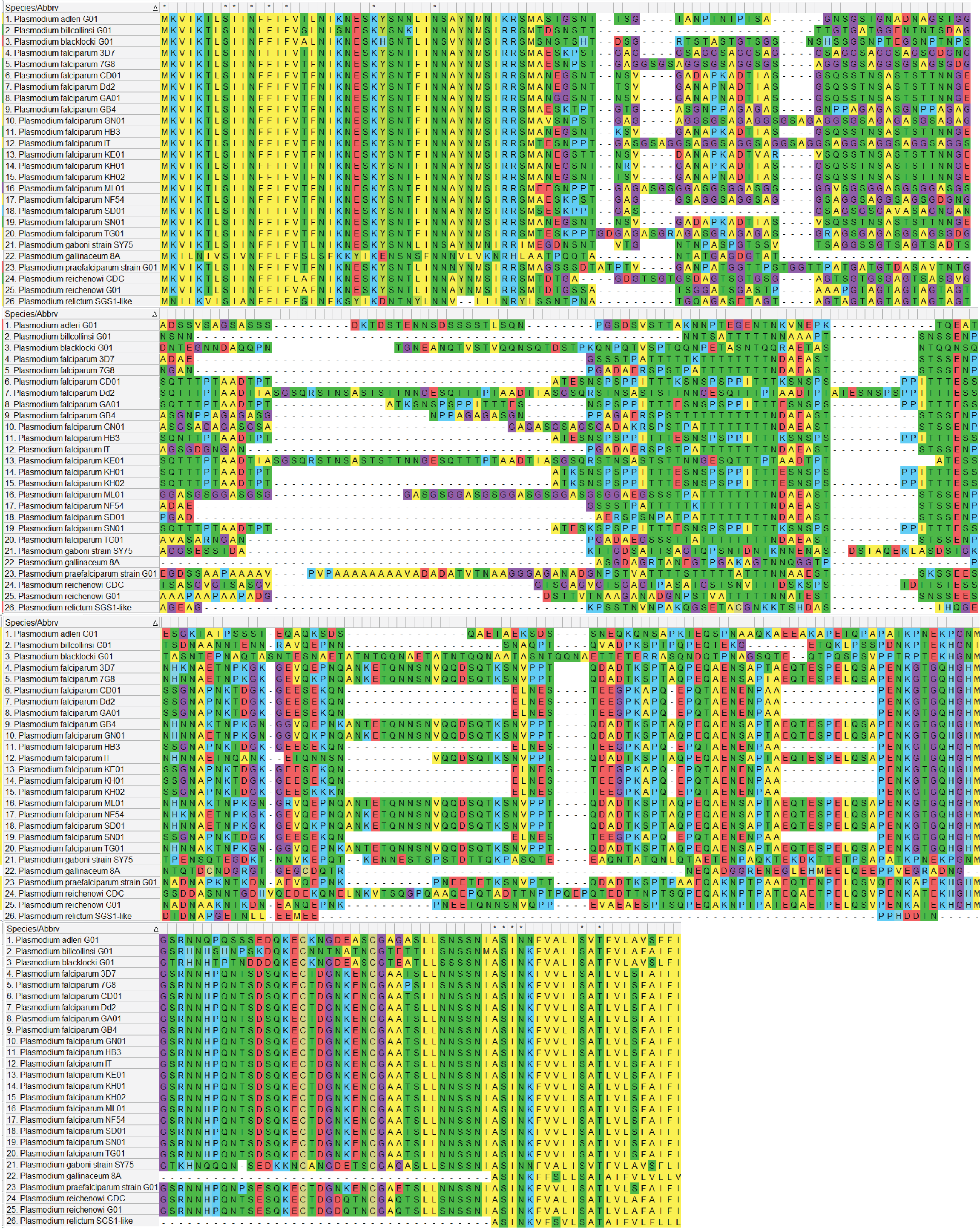

**Supplementary Figure 1** Alignment of full-length amino acid protein sequences for all *Pf*MSP2 like sequences on PlasmoDB. Alignment done using MUSCLE alignment with default settings.

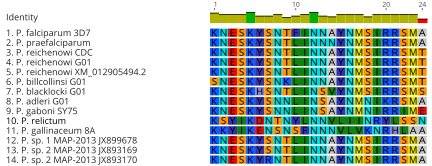

**Supplementary Figure 2** Alignment (Blosum 62) of Amino acid sequences for the N-terminal conserved region of *P. falciparum* 3D7, available *Laverania*, *P. relictum* and *P. gallinaceum* MSP2 like sequences on PlasmoDB and identified through a BLAST search on the NCBI server. *Plasmodium*. sp. 1, 2 and 3 represent MSP2 sequences identified using a BLAST search that are indicated to be from an unspecified *Laverania* spp. parasite. Amino acids are coloured according to their properties (RasMol).

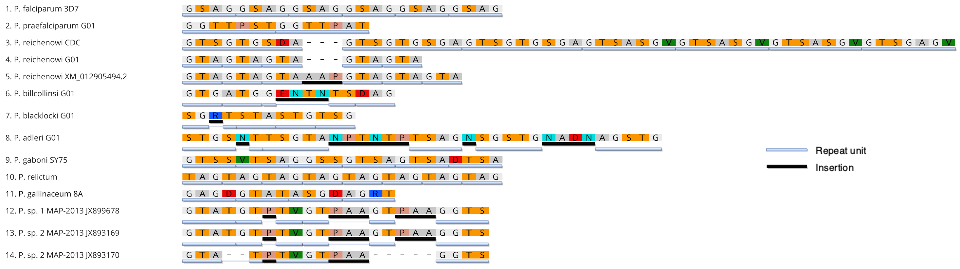
**Supplementary Figure 3** Amino Acid sequences for the N-terminal repeat region of *P. falciparum* 3D7, available *Laverania*, *P. relictum* and *P. gallinaceum* MSP2 like sequences on PlasmoDB and identified through a BLAST search on the NCBI server. *Plasmodium*. sp. 1, 2 and 3 represent MSP2 sequences identified using a BLAST search that are indicated to be from an unspecific *Laverania* spp. parasite. Sequences presented are not aligned, with the exceptions of the 3 *P. reichenowi* and 3 unspecified *Laverania* sequences in order to highlight the conserved sequence despite an insertion. Insertions that are considered to break up the small amino acid and/or repeat structure for the purposes of this comparison are underlined in Black. Respeats, both conserved and degenerate, are underlined in grey. Amino acids are coloured according to their properties (RasMol).

**Supplementary Figure 4** Alignment of amino acid sequences for the region corresponding to the C-terminal conserved region of *P. falciparum* 3D7, available *Laverania*, *P. relictum* and *P. gallinaceum* MSP2 like sequences on PlasmoDB and identified through a BLAST search on the NCBI server. *Plasmodium. sp.* 1, 2 and 3 represent MSP2 sequences identified using a BLAST search that are indicated to be from an unspecified *Laverania* spp. parasite. Initial alignment done using Blosum62 with default settings, with subsequent manual modification as required to align to the cysteine residues and other prominent areas of conservation between the *Laverania* and bird malarias. Amino acids are coloured according to their properties (RasMol).

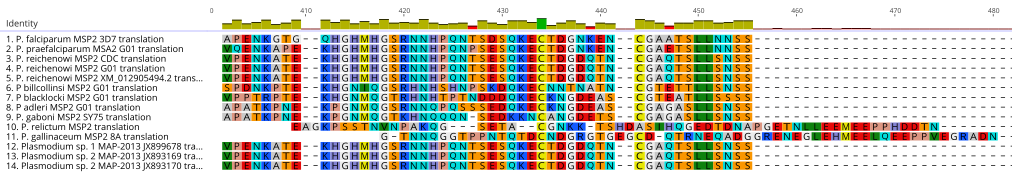

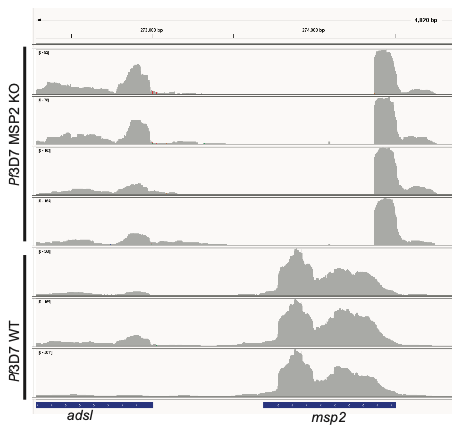

**Supplementary Figure 5** Coverage plot showing absence of the majority of *msp2* in the schizont transcriptome of *Pf*3D7 ΔMSP2 Clone 1 parasites but *msp2* presence in *Pf*3D7 MSP2 WT. Small section of *msp2* transcribed in *Pf*3D7 ΔMSP2 Clone 1 parasites corresponds with the end of the homology region and primarily covers the signal peptide.
