## Supplementary Tables 1 to 4 for "An abundant merozoite surface protein of *Plasmodium falciparum* modulates susceptibility to inhibitory antibodies"

**Supplementary Table 1.** Primers used to generate MSP2 KO lines.

| | | |
|---|---|---|
| 5' flank MSP2 KO F | GGTCCGCGGCTCGACTAATCAATTTACAATTTC | Sac11 |
| 5' flank MSP2 KO R | GGTACTAGTGCCATACTTCTCCTTATACTC | Spe1 |
| 3' flank MSP2 KO F | GGTGAATTCCTCTTCATTTTAAAACATTGAC | Ecor1 |
| 3' flank MSP2 KO R | GGTCCATGGGTTTTTTCAATGCGTGC | Nco1 |
| Dd2 MSP2 KO 5' flank F | GGTCCGCGGCTCGACTAATCAATTTACAATTTCC | Sac11 |
| Dd2 MSP2 KO 5' flank R | GGTACTAGTCTAGTAGTATTAGAACCTTCATTTG | Spe1 |
| Dd2 MSP2 KO 3' flank F | GGTGAATTCATTCTCTTCATTTTAAAACATTGAC | EcoR1 |
| Dd2 MSP2 KO 3' flank R | GGTCCATGGGTACTTGAAGAAATATGGTACC | Nco1 |
| PfMSP2 guide 1 F | TAAGTATATAATATTTGGTAATGGTGCAGATGCTGGTTTT AGAGCTAGAA | |
| PfMSP2 guide 1 R | TTCTAGCTCTAAAACCAGCATCTGCACCATTACCAAATATT ATATACTTA | |
| Dd2 MSP2 Guide 2 F | TAAGTATATAATATTGCACCAGAGAATAAAGGTACGTTTT AGAGCTAGAA | |
| Dd2 MSP2 Guide 2 R | TTCTAGCTCTAAAACGTACCTTTATTCTCTGGTGCAATAT TATATACTTA | |
| 5' int 3D7 MSP2 KO F | CTTTTCATATAATAAATCAAATGCAC | |
| 5' int 3D7 MSP2 KO R | TACAAAATGCTTAAGACAGATC | |
| 3D7 WT 5' int check R | GGTAGTTGTGGTAGTAGC | |
| Dd2 5' check F | CAATTACGATATAAAACCTAGTATCTTTC | |
| Dd2 WT 5' int check R | CTATTTGTACTCCTTTGACTTCCAC | |

**Supplementary Table 2.** Antibodies used in this study.

| Serum/Antibody/ibody | Source |
|---|---|
| Anti-MSP2 mAb 2F2 | Adda et al Infect Immun 2012 |
| Anti-MSP2 Rb FC27 | Robin Anders, LaTrobe University |
| Anti-EXP2 Rb | Bullen et al. J Biol Chem 2012 |
| Anti-PfAldolase | Abcam |
| Anti-PfRh2b Rb 1055-3 | Lopaticki et al Infect and Immun 2011 |
| Anti-PfEBA175/Rh2b Rb 1056-3 | Lopaticki et al Infect and Immun 2011 |
| Anti-PfEBA175/Rh4 Rb 1049-3 | Lopaticki et al Infect and Immun 2011 |
| Anti-PfEBA175/Rh2b/Rh4 Rb 1067-3 | Lopaticki et al Infect and Immun 2011 |
| Anti-w2mef EBA175 Rb | Healer et al PLoS One 2013 |
| Anti-MSP1-19 Rb (6858, 645, 647) | Paul Gilson, Burnet Institute |
| Anti-Rh5.1 Rt 494 | Healer et al Front Cell Infect Microbiol 2022 |
| Anti-PTRAMP nAb H8 | Scally et al Nat Microbiol 2022 |
| Anti-CSS nAb D2 | Scally et al Nat Microbiol 2022 |
| Anti-CSS nAb D2-FC | Scally et al Nat Microbiol 2022 |
| Anti-AMA1 3D7 Rb (1072, 1149, 1161) | Drew et al. PLoS One 2012 |
| Anti-AMA1 w2mef Rb (1151, 1164, 1163, 128/7) | Drew et al. PLoS One 2012 |
| Anti-AMA1 mAb 1F9 | Coley et al PEDS 2001 |
| Anti-AMA1 mAb 4G2 | Collins et al J. Biol. Chem. 2007 |
| Anti-AMA1 ibody WD33 | Angange et al Under Review |
| Anti-AMA1 ibody WD34 | Angange et al Under Review |
| Anti-AMA1 WD34 | Angange et al Under Review |
| Anti-AMA1 ibody WD34-FC | Angange et al Under Review |

Rb= rabbit. mAb= mouse monoclonal. Rt= rat. nAb= camelid nanobody. I-body= single domain antibody derived from shark variable new antigen receptor ($V_{NAR}$).

**Supplementary Table 3.** Enzymes used to remove/inactivate RBC receptors and target residues of the enzyme.

| **Enzyme** | **Cleavage Target** | **Supplier** |
|---|---|---|
| Neuraminidase (0.067 U/mL) | Sialic acid residues | Sigma-Aldrich |
| High Trypsin (1 mg/mL) | Lysine or Arginine residues, unless followed by a proline | Sigma-Aldrich |
| Low Trypsin (0.067 mg/mL) | Lysine or Arginine residues, unless followed by a proline | Sigma-Aldrich |
| Chymotrypsin (1 mg/mL) | Tyrosine, Tryptophan or Phenylalanine | Worthington Biochemical Corporation |
| Chymotrypsin (1 mg/mL)/ Trypsin (0.067 mg/mL) | Tyrosine, Tryptophan, Phenylalanine, Lysine or Arginine residues, unless followed by a proline | Worthington Biochemical Corporation & Sigma-Aldrich |

**Supplementary Table 4.** Primers used for RT- qPCR of *P. falciparum* MSP2, MSP5, MSP4 and controls.

| Name | Sequence |
|---|---|
| FrucBiAld 3D7 F qPCR [a, b] | TGTACCACCAGCCTTACCAG |
| FrucBiAld 3D7 R qPCR [a,b] | TTCCTTGCCATGTGTTCAAT |
| MSP2 3D7 F qPCR | TCCTACTGCACAACCTGAACAA |
| MSP2 3D7 R qPCR | ATGTCCATGTTGTCCTGTACCTT |
| MSP4 3D7 F qPCR | CAAAAGAATCCCAAATGGTTGATGATAA |
| MSP4 3D7 R qPCR | AACATGGCCACCTGAATTTGAT |
| MSP5 3D7 F qPCR | AAATTAATGAGAATGCAGAAATAGGTCAA |
| MSP5 3D7 R qPCR | CGCTATAATGTGGCACCTCAT |
| SUB1 3D7 F qPCR[c] | GGAATGAGGTAGATGCCGATGAA |
| SUB1 3D7 F qPCR[c] | TCCTTTACATCTTTCAGTTCCCTCAT |

[a] FrucBisAld primers had been previously validated as an appropriate housekeeping gene by Salanti et. al. (2003). Primers listed here are the fructose-bisphosphate aldolase qPCR primers (primer pair 61) from Salanti et. al. (2003).

[b] fructose-bisphosphate aldolase is ubiquitously expressed throughout the parasite lifecycle and is used as a non-stage specific housekeeping gene.

[c] SUB1 is a schizont expressed protein and is used as a schizont-stage specific housekeeping gene.
